# Gastric-Brain Coupling Dynamics in Response to a Cognitive Challenge Reveal Distinct Autonomic and Psychological Correlates in Healthy Individuals and Patients with Irritable Bowel Syndrome Gastric-Brain Coupling Psychophysiological Correlates

**DOI:** 10.64898/2026.07.21.739524

**Authors:** Jeanne Rudy, Minjoz Séphora, Bonaz Bruno, Sinniger Valérie, Hot Pascal, Kibleur Astrid, Pellissier Sonia

**Author notes:** Corresponding author : Rudy JEANNE. Co-first authors (Jeanne Rudy, Minjoz Séphora). Authors who contributed equally as co-supervisors of the project (Hot Pascal, Kibleur Astrid, Pellissier Sonia).

## Abstract

The gastric’s intrinsic slow-wave rhythm couples to ongoing cortical activity in humans, but the functional significance of this gastric-brain coupling remains poorly understood. Using simultaneous electroencephalography and electrogastrography, we examined how gastric-brain coupling changes during a mental task and subsequent recovery and whether these dynamics relate to autonomic responses in healthy adults (HA) and individuals with Irritable Bowel Syndrome (IBS), a disorder of gut-brain interactions. In HA, gastric-brain coupling decreased during mental task and gradually returned to baseline during recovery, primarily for cortical alpha rhythms. Greater reductions in gastric-brain coupling were associated with stronger sympathetic arousal, indicating that gastric-brain coupling is associated with physiological responses to cognitive demand. In IBS, gastric-brain coupling dynamics were preserved. However, unlike HA, sympathetic arousal was not associated with gastric-brain coupling reductions, suggesting an alteration in the functional relationship between gastric-brain coupling and autonomic regulation despite preserved coupling dynamics. Further, in HA, greater reductions in gastric-brain coupling were associated with higher anxiety and neuroticism and lower interoceptive awareness. In contrast, these relationships were reversed in IBS, with greater gastric-brain coupling dynamics observed in individuals with higher anxiety and neuroticism and lower interoceptive awareness. Together, gastric-brain coupling emerges as a dynamic physiological process associated with autonomic responses to cognitive demand. In IBS, these dynamics are preserved but their physiological and psychological associations are altered, indicating a dissociation between gastric-brain coupling and its functional correlates.

**Significance Statement:** The functional role of gastric-brain coupling remains largely unknown. We show that coupling between the gastric’s intrinsic slow-wave rhythm and cortical activity is dynamically reduced during mental engagement and recovers afterward, primarily through alpha rhythms. In healthy adults, these changes follow sympathetic arousal, linking gastric-brain coupling to autonomic responses to cognitive demands. In irritable bowel syndrome, coupling dynamics remain intact but lose their relationship with autonomic responses and show altered associations with anxiety, neuroticism, and interoceptive awareness. These findings suggest that gastric-brain coupling contributes to the physiological organization of brain-body states and that irritable bowel syndrome may involve altered functional interpretation of visceral signals despite preserved gastric-brain coupling dynamics.

## 1. Introduction

Recent evidence has revealed a novel form of gastric-brain interaction in humans. Gastric slow waves, electrical oscillations generated by the interstitial cells of Cajal at approximately 0.05 Hz, synchronize with cortical alpha oscillations (Richter et al., 2017) and resting-state bold signal oscillations across sensory, motor and associative regions (Rebollo et al., 2018; Rebollo & Tallon-Baudry, 2022). These findings suggest that gastric rhythm may contribute to large-scale brain dynamics (Azzalini et al., 2019; Engelen et al., 2023; Rebollo et al., 2021).

Despite increasing support for the existence of gastric-brain coupling, its functional significance remains poorly understood (Porciello et al., 2018; Rebollo et al., 2021). A central unresolved question is whether this coupling merely reflects a stable physiological background phenomenon observable at rest or whether it is dynamically associated with physiological reactivity. Only a few pilot studies have examined the sensitivity of gastric-brain coupling to changing psychophysiological states, yielding mixed results. For example, Balasubramani et al. (2022) reported reduced frontal gastric-brain coupling during cognitive task performance compared with rest. In contrast, our previous work showed reduced coupling during post-task cardiorespiratory relaxation (Jeanne et al., 2023). Furthermore, because most of the previous studies focused exclusively on alpha oscillations it remains unclear whether gastric-brain coupling extend to faster cortical rhythms associated with executive processing (Jensen & Colgin, 2007; Keitel et al., 2022; Lundqvist et al., 2024).

One possibility is that gastric-brain coupling reflects a dynamic mechanism supporting physiological information exchanges between these two organs (Engelen et al., 2023). Gastric activity is monitored and regulated by the central nervous system through autonomic nervous system (ANS) pathways (Holtmann & Talley, 2014). This regulation involves bidirectional interactions between the stomach and the central nervous system that are highly sensitive to cognitive and emotional states (Critchley & Harrison, 2013; Holtmann & Talley, 2014; Porciello et al., 2024). If gastric-brain coupling contributes to the integration of visceral processes, its strength should vary with cognitive demands and be associated with autonomic responses. Specifically, cognitive challenge and subsequent recovery are expected to elicit coordinated sympathetic and parasympathetic adjustments that can be assessed using electrodermal activity (EDA; Critchley, 2002; Sequeira et al., 2009) and heart rate variability (HRV) measures (Laborde et al., 2017, 2018). Within this framework, task-related changes in gastric-brain coupling should be associated with autonomic responses, supporting a role for gastric-brain coupling in adaptive viscera-brain regulation.

Examining gastric-brain coupling dynamics in both healthy individuals and patients with Irritable Bowel Syndrome (IBS) can reveal whether it represents a general adaptative mechanism and whether it can be disrupted in a disorder of gut-brain interaction. IBS is particularly relevant in this context because it is characterized by several alterations involving neurovisceral integration such as heightened visceral sensitivity, exacerbation of gastrointestinal symptoms under stress, altered autonomic regulation, and disturbances in interoceptive processing (Drossman & Hasler, 2016; Enck et al., 2016; Lacy et al., 2021; Mayer et al., 2023; Pellissier & Bonaz, 2017; Sadowski et al., 2020; Ying-Chih et al., 2020). If gastric-brain coupling contributes to adaptive brain-body regulation, alterations in its temporal dynamics and physiological associations would be expected in IBS relative to healthy individuals.

The first objective of the present study was to determine whether phase-amplitude gastric-brain coupling exists across cortical rhythms and how it dynamically changes during and immediately after a cognitive challenge. The second objective was to evaluate whether coupling dynamics covary with sympathetic and parasympathetic responses indexed by EDA and HRV. We further examined whether coupling dynamics and their autonomic correlates differ between IBS participants and healthy adults (HA). Finally, as an exploratory objective, we examined whether gastric-brain coupling dynamics are associated with interoceptive awareness and emotional regulation tendencies.

## 2. Materials and Methods

Some parts of the methods were not processed or analyzed in this article, as they are part of a larger project that includes two different empirical studies focused on heart rate variability biofeedback (HRV-BFB) effects on psychophysiology published elsewhere (Minjoz et al., 2025, 2026). However, we report the full experimental design performed on participants, but provide details only for those described herein.

### 2.1. Participants and Ethics Statement

This study includes data from two complementary studies: a clinical trial involving adults with IBS, and a randomized controlled trial with HA. The first trial was approved by the national research ethics committee of SUD-EST VI Clermont-Ferrand, France (reference AU 1658; 2020-A02155-34) and registered in ClinicalTrials.gov (Identifier: NCT04807933). The second was approved by the ethics committee of University of Savoie Mont Blanc, France (reference C.E.R.E.U.S: CER_2021_10) and pre-registered on OSF (https://doi.org/10.17605/OSF.IO/FKB79). All participants provided written informed consent. Participants eligible for inclusion were 18-70 yrs old and not experts in relaxation practices. Exclusion criteria were a diagnosis of neurological, psychiatric, cardiac diseases; taking any medication that could influence the ANS (e.g., α- or β-blockers); and being pregnant. Participants who were included were asked not to consume stimulating drinks or to use tobacco 2 hrs before the experiment and not to undergo physical activity 12 hrs before the experiment (Laborde et al., 2017). They were also asked to eat a light breakfast at least 1 hr before the session, ensuring a minimum of 2 hrs had elapsed before the start of the electrogastrogram (EGG) recordings (Yin & Chen, 2013). Additionally, participants were instructed not to wash their hair the day before the session for electroencephalogram (EEG) requirements.

Sample size was computed *a priori* based on the main effect of cardiac responses to HRV-BFB practice (Minjoz et al., 2025; Minjoz et al., 2026). The sample size is similar or greater than the most recent studies on EGG-EEG gastric-brain coupling (Balasubramani et al., 2022; Jeanne et al., 2023; Richter et al., 2017). Sixty-two HA (49 females; M*_age_* = 36.13 yrs; SD*_age_* = 14.76 yrs) were recruited between March 2021 and August 2022 from a call for participation in a psychology study and received 50€ in compensation. The highest level of education attained by the participants was a university degree (53.23%), followed by a high school diploma (32.26%) and a general certificate of secondary education (6.45%). Five participants were retired (8.06%). Eight participants discontinued the protocol prior to completion and were excluded from analysis. An additional nine HA were excluded due to poor signal quality based on predefined EGG and EEG artifact rejection criteria (see Physiological Measurements). As a result, 52 HA were included in Session 1 and 53 in Session 2. A final subsample of 45 HA (34 females; M*_age_* = 36.4 yrs; SD*_age_* = 15.1 yrs) had analyzable data across both sessions and were included in the analyses.

Thirty-three adults with IBS (23 females; M*_age_* = 40.61 yrs; SD*_age_* = 13.30 yrs; 13 mixed IBS, 2 IBS with constipation, and 18 IBS with diarrhea) were recruited between March 2021 and March 2023 from the division of Hepato-Gastroenterology of Grenoble University Hospital and met Rome IV diagnostic criteria (Drossman & Hasler, 2016). Comorbidities in this group included fibromyalgia (n = 3), endometriosis (n = 4), and attention-deficit disorder (n = 1). Average IBS diagnostic delay was 4.2 yrs (SD = 6.3 yrs), with symptom onset reported at an average of 12.4 yrs (SD = 11.2 yrs) prior the study. The highest level of education attained by the participants was a university degree (54.55%), followed by a high school diploma (36.36%) and a general certificate of secondary education (9.09%). Four IBS discontinued the protocol prior to completion and were excluded from analysis. An additional five IBS were excluded due to poor signal quality based on predefined EGG and EEG artifact rejection criteria (see Physiological Measurements). A subsample of 24 adults with IBS (18 females; M*_age_* = 40.8 yrs; SD*_age_* = 13.2 yrs) had usable EEG and EGG data at both sessions and were included in the analyses.

### 2.2. Procedure

Participants completed three experimental sessions, each separated by a 24-day period (Figure 1). Participants did not practice any training during the first period and had to practice relaxation training (i.e., HRV-BFB) daily during the second period (the latter is not analyzed in the current study, for details see Minjoz et al., 2026). Participants were asked to complete psychological questionnaires between each session (see Psychological Questionnaire section). The three sessions occurred between 9:00 AM and 12:00 PM in a temperature-controlled room (23°C). In all sessions, participants completed self-report psychological state questionnaires (not analyzed in this study). Then electrophysiological sensors were placed (EEG, EGG, electrooculogram [EOG], electrocardiogram [ECG], EDA, photoplethysmogram [PPG, not analyzed] and respiratory belt [not analyzed]). Then participants sat comfortably in a medical reclining chair for a 30-min eyes-open resting state (i.e., rest module). This module consisted in staring at a cross on a computer screen, remaining silent and still, letting their mind wander and avoid excessive movement. Following this, a 10-min computerized mental task (i.e., task module) was proposed. This was a modified Stroop task (coded in OpenSesame) which involved color-word congruency judgments under increasing time pressure and cognitive load across six mini-blocks. Every two blocks, response keys were reversed, and a concurrent working-memory task (color count) was added in the final blocks. Visual feedback on performance, a countdown timer, and a fictitious social comparison metric (+10% above actual performance) were included to increase mental load (for detailed explanations about this task, see Minjoz et al., 2026). Then, a final 30-min eyes-open resting state (i.e., recovery module) was done. Perceived stress ratings (visual analog scales [VAS]) were also collected at rest, after the task and after the recovery modules.

**Figure 1:**
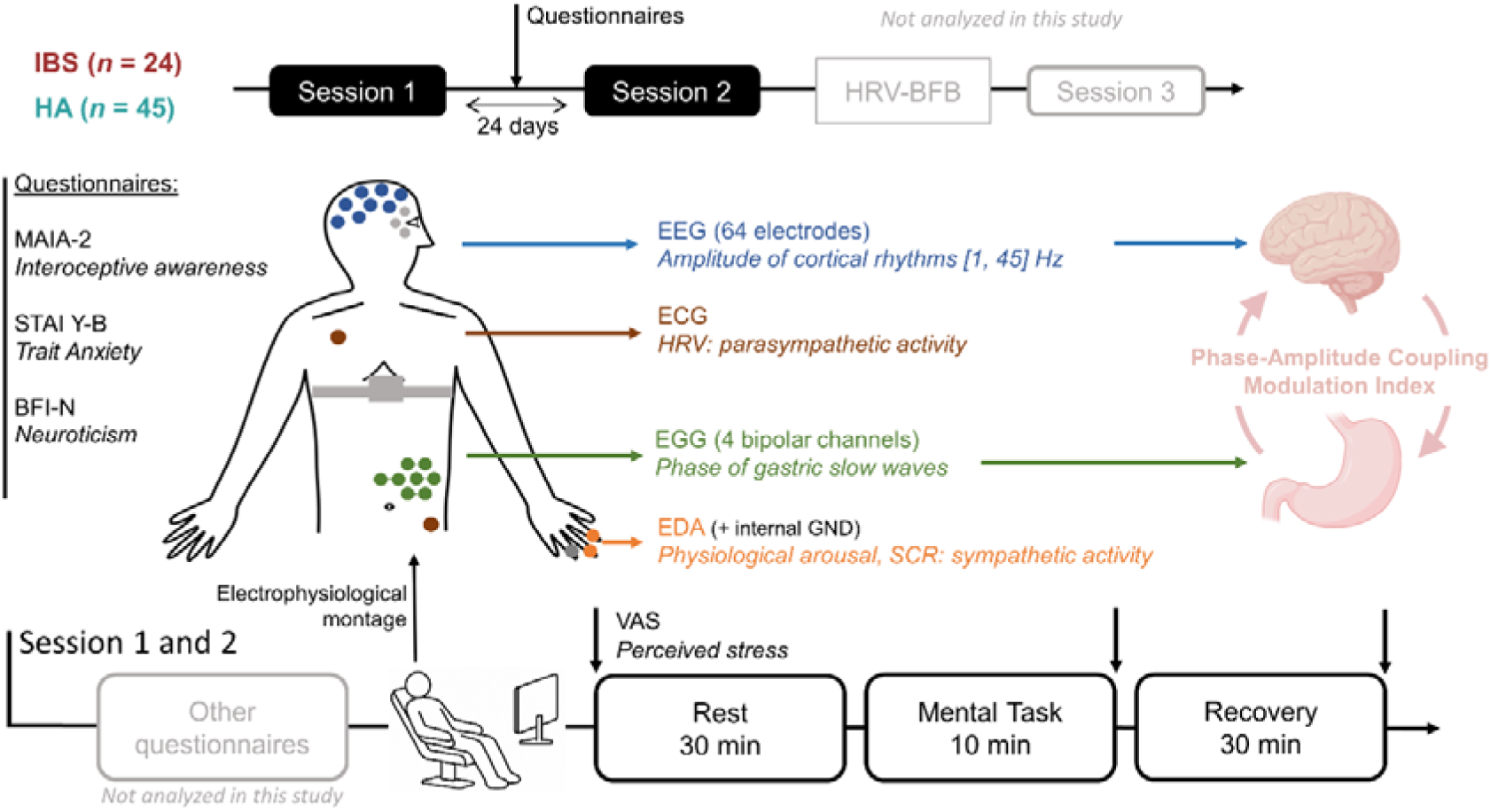
Graphical representation of the protocol design. The current study does not include the evaluation of the heart rate variability biofeedback. IBS: Irritable Bowel Syndrome, HA: Healthy Adults. VAS: Visual Analog Scale. EEG: Electroencephalogram, ECG: Electrocardiogram, EGG: Electrogastrogram, EDA: ElectroDermal Activity. GND: Ground. HRV-BFB: Heart Rate Variability Biofeedback. Pictograms were drawn from BioRender.

### 2.3. Psychological Measurements

#### 2.3.1. Questionnaires

The sensitivity to experiencing negative emotion was evaluated by the 5-item neuroticism subscale of the Big Five Inventory (BFI-N, John et al., 1991; French version, Plaisant et al., 2010). The trait anxiety was evaluated using the 20-item Trait Subscale of State-Trait Anxiety Inventory (STAI Y-B, Spielberger et al., 1983; French version, Spielberger et al., 1993). Interoceptive awareness was evaluated with the 37-item Multidimensional Assessment of Interoceptive Awareness version 2 (MAIA-2, Mehling et al., 2018; French version, Willem et al., 2022). It is composed of eight dimensions: (1) Noticing (i.e., awareness of uncomfortable, comfortable, and neutral bodily sensations); (2) Not-Distracting (i.e., tendency to ignore sensations of discomfort or pain); (3) Not-Worrying (i.e., tendency not to worry about, or, experience emotional distress related to discomfort or pain); (4) Attention Regulation (i.e., ability to sustain and control attention toward bodily sensations); (5) Emotional Awareness (i.e., awareness of the connection between bodily sensations and emotional states); (6) Self-Regulation (i.e., ability to regulate distress through attention to bodily sensations); (7) Body Listening (i.e., active listening to the body to better understand internal states); (8) Trusting (i.e., trust in bodily sensations). Perceived stress was evaluated using the average score of four VAS ranging from 0 (not at all) to 10 (extremely) with the following questions: “How nervous do I feel?”, “To what extent are things going my way?” (reverse-scored), “How relaxed do I feel?” (reverse-scored), and “How calm do I feel?” (reverse-scored). Due to technical issues, data from 38 HA and 23 IBS were analyzable regarding the perceived stress.

#### 2.3.2. Analysis

Perceived stress was analyzed using linear mixed-effects models, implemented in R with the lme4 package. Fixed effects included modules (i.e., rest, task, recovery), sessions and groups (all sum contrasted) and their interactions. Only random intercepts were modeled for participants after choosing the parsimonious, converging and non-singular mixed model following Bates methodology (Bates et al., 2015). Perceived stress was square root transformed to fulfill linear mixed modelling assumptions (i.e., normality, linearity, homoscedasticity), visually checked using the performance package (Lüdecke et al., 2021). Model comparisons were conducted using likelihood ratio tests via the type 3 anova function. Post-hoc pairwise contrasts were computed for the effect of module and corrected using Tukey’s method, and, polynomial contrasts for the module × group interaction using the emmeans package (Lenth & Piaskowski, 2026).

### 2.4. Physiological Measurements

#### 2.4.1. Acquisition and Preprocessing

EEG signals were collected by using Active-Two amplifier from BioSemi at a sampling frequency of 1024 Hz. Peripheral signals (i.e., EGG, ECG, EDA, PPG, respiration) were collected by using BIOPAC (MP 150, CEROM, France) at a sampling frequency of 1000 Hz.

##### 2.4.1.1. Electroencephalogram

EEG signals were recorded using a 64-channel active electrode cap (10-20 standard layout) at a bandwidth of 0-208 Hz. Data preprocessing was conducted using the MNE-Python library (Gramfort, 2013). Bad EEG channels (i.e., channels delivering no signal or exhibiting extreme variability due to poor scalp contact) were excluded from the analysis. A band-pass FIR filter between 1-40 Hz and a notch FIR filter between 48-52 Hz (with default MNE filters settings) were applied to reduce the 50 Hz power line noise and isolate relevant EEG rhythms. Data were then segmented into 2-second-long epochs and downsampled at 200 Hz to reduce computational load. Independent Component Analysis (ICA) using the fastICA algorithm (Ablin et al., 2018) was performed to remove ocular, cardiac and muscular artifacts components. The AutoReject algorithm (Jas et al., 2017) was applied using default parameters (n_interpol = [4, 8, 12]; consensus = {i × 0.1, i ∈ [1:10]}) to automatically reject epochs containing high-variance artifacts (e.g., transient electrode disconnections) and to interpolate noisy electrodes within individual epochs. The preprocessed EEG data were visually inspected, and any remaining noisy epochs were manually excluded. Finally, removed EEG channels were interpolated (using spline interpolation) and EEG channels were referenced to the common average across all electrodes. Twenty Hz sampled EEG power time series per EEG electrodes and per module was extracted by taking the squared Fourier transform coefficient from 1 to 45 Hz at 1Hz step in 95% overlapping one-second-long sliding Hanning windows (Richter et al., 2017).

##### 2.4.1.2. Electrogastrogram

Gastric slow waves were recorded using four single-use bipolar cutaneous Ag/AgCl electrodes and placed on the abdominal skin previously cleaned with alcohol and exfoliated with an abrasive gel. All electrodes were 4 cm spaced with the negative electrode closer to the umbilicus than the positive electrode in the bipolar montage and followed the standardized montage illustrated in Figure 1 and described in (Wolpert et al., 2020). EGG was acquired using EGG100C BIOPAC bioamplifier at a bandwidth of 0 -1 Hz and a 1000 amplification gain and then downsampled to 20 Hz. In accordance with current literature recommendations (Wolpert et al., 2020), the gastric slow waves were extracted for the bipolar electrodes using the sharpest and highest spectral power peak in the normogastric frequency range (2-4 cycles per min). To do so, the Welch power spectrum of the EGG signals per experimental session was derived using a 200-second-long window with 75% overlap. Participants were not further analyzed in case of absence of a clear normogastric peak. The EGG signal was then filtered using a FIR filter around the frequency of the normogastric peak for the selected EGG electrode (order = 3 x sampling frequency/low frequency limit of filter) ± 0.015 Hz. Filtered EGG recordings containing cycles of abnormally long or short duration and a non-monotonic phase, were removed from the analyses, following Wolpert and colleagues’ methodology. No significant difference in proportion of deleted cycles between modules was observed (aligned rank transform ANOVA; *F*(4, 612) = 2.09, *p* = .08). Normogastric power, mean and standard deviation of normogastric cycle durations were extracted as indicators of gastric slow waves dynamics per module.

##### 2.4.1.3. Electrocardiogram

ECG was recorded using two single-use cutaneous Ag/AgCl electrodes placed on the right shoulder and left subcostal area (Figure 1). ECG was acquired at a bandwidth of 0.05-35 Hz and a 1000 amplification gain using ECG100C BIOPAC bioamplifier. ECG signal visualization and R peak detections were performed using the NeuroKit2 Python library with custom scripts replicating Kubios software processing parameters as well as its automatic artifact correction (Lipponen & Tarvainen, 2019). The pre-processed R peaks detected were used to obtain the interbeat interval time series and derived the HRV signals. The root mean squared of HRV successive differences (HRV-RMSSD) was extracted per module as an index of parasympathetic activity (Laborde et al., 2017) using NeuroKit2.

##### 2.4.1.4. Electrodermal Activity

EDA was recorded using skin conductance sensors (SS3LA, BIOPAC) filled with isotonic gel and placed on the third phalanx of the left index and middle fingers. EDA was acquired at a bandwidth of 0-10 Hz and a 10 µS/V amplification gain using GSR100C BIOPAC bioamplifier. The signal was then bandpass filtered between 0 and 3 Hz and downsampled to 20 Hz. Tonic and phasic components were extracted using the cvxEDA algorithm (Greco et al., 2016). Nonspecific skin conductance responses (EDA-SCR) were defined when the phasic peak of electrodermal responses exceeded 0.05 µS (Boucsein et al., 2012). The mean amplitude of EDA-SCR was calculated using NeuroKit2’s as an indicator of physiological arousal and descending cholinergic sympathetic influences upon sweat glands (Critchley, 2002; Sequeira et al., 2009). When no EDA-SCR were detected, the EDA-SCR mean was set to 0 µS.

#### 2.4.2. Analyses

From the 30 min rest module, only the 10-20 min segment was retained for analyses to reduce anxiety-related artifacts from electrode placement, keeping the most engaging period. The recovery period was divided into three 10-min-long segments. The autonomic and gastric slow waves indices were analyzed using linear mixed-effects models and following the same methodology as perceived stress (see paragraph 2.3.2). If necessary, power transformations have been used to fulfill the modelling assumptions. Visualizations of linear mixed modelling effects were generated using the ggeffect package (Lüdecke, 2018).

### 2.5. Gastric-Brain Coupling

#### 2.5.1. Extraction

Gastric-brain phase-amplitude coupling was quantified using the modulation index (MI) following the method described by Tort and colleagues (2010). Briefly, the phase of the preprocessed EGG signal was extracted using the Hilbert transform and divided into 18 equal bins. Then, the EEG power time series was averaged per EGG phase bins for each frequency (1-45 Hz) and electrode per module. This provides a phase-amplitude profile where a strong EGG-phase EEG-amplitude coupling would be reflected in an important variation of EEG amplitude across the EGG phase bins. The MI was then computed by quantifying the Kullback-Leibler divergence between the observed phase-power distribution and a uniform distribution. The resulting MI tends to 1 in case of high phase-amplitude coupling and tends to 0 in case of an absence of coupling. However, because of natural oscillations of the gastric pace-setter rhythm and cortical activity, even in the absence of significant phase-amplitude coupling, the MI will not be equal to 0. Therefore, an appropriate chance-level MI needs to be determined to test the significance of the MI and avoid spurious detection of phase-amplitude coupling (Aru et al., 2015). The chance-level MI has been computed per participant, frequency, channel and experimental session as the median of 1000 surrogates MI computed by circularly shifting the EGG phase by at least 60 s (around 3 normogastric cycles) compared to the EEG power time series. The strength of the gastric-brain coupling is calculated as the difference between the real MI and the chance level per participant, experimental session, EEG channel and frequency.

#### 2.5.2. Analysis

The study of gastric-brain coupling dynamics followed two steps. The first step consisted in identifying the electrodes and frequencies of EEG power significantly coupled to the phase of the stomach pace-setter (compared to the chance-level MI) in the entire experimental sessions. It was tested using cluster-based spatio-spectral permutation testing (Maris & Oostenveld, 2007) per step of 1 Hz including all EEG channels. Then the gastric-brain coupling was averaged in delta [1, 4[ Hz, theta [4, 8[ Hz, alpha [8, 13[ Hz, beta [13, 30[ Hz and gamma [30, 45[ Hz rhythms for HA and IBS and cluster-based spatial permutation testing was done per cortical rhythm and group. IBS were compared with HA at each cortical rhythm using cluster-based spatial permutation of two sampled Welch t-tests statistics. The threshold for cluster inclusion was set to α = .05 and 5000 permutations were performed.

The second step consisted in testing the effect of the modules and sessions on gastric-brain coupling. To do so, only the frequency and electrodes included in a significant cluster of existing gastric-brain coupling along the session were further processed. From the 30 min rest module, only the 10-20 min segment was retained for analyses to reduce anxiety-related artifacts from electrode placement, keep the most engaging period and get the same baseline duration than task duration as the MI is sensitive to data length (Tort et al., 2010). The recovery period was divided into three 10-min-long segments. A cluster-based spatio-spectral permutation testing of a 5 (modules) × 2 (sessions) repeated-measures ANOVA was conducted on gastric-brain coupling strength to detect module- and session-related effects for HA. IBS were compared with HA at each module using cluster-based spatial permutation of two sampled t-tests between groups on gastric-brain coupling averaged in the 10-13 Hz interval where the effect of modules is the most important and robust. For each group, post-hoc pairwise module comparisons were performed using cluster-based spatial permutation tests. Cluster p-values were corrected for multiple comparisons using Holm-Bonferroni method (Holm, 1979). For each of these analyses, the threshold for cluster inclusion was set to α = .05 and 5000 permutations were performed.

### 2.6. Psychological Correlates of Gastric-Brain Coupling Dynamics

To examine how gastric-brain coupling at 10-13 Hz significant modules differences covaried with significant module differences on perceived stress, EDA-SCR and HRV-RMSSD, we performed cluster-based spatial permutation testing on Spearman correlation statistic (MNE-Python; 5000 permutations, cluster-forming threshold set to alpha = .05).

Gastric-brain alpha coupling exhibited a quadratic dynamic across the scalp, both in IBS and HA. For each participant, gastric-brain coupling was averaged in part of the alpha band (10-13 Hz) for each electrode and across the five 10-min modules within rest, task, and recovery, during both experimental sessions. The windows were labeled 0, 20, 30, 40, 50, corresponding to the midpoint of each time bin, with minute 15 of rest set as time zero. For each electrode and participant, second-degree polynomial coefficients modeling gastric-brain coupling dynamics were computed using the polyfit function from the NumPy Python package. To assess whether MAIA-2, STAI Y-B, and BFI-N were associated with gastric-brain coupling dynamics in HA, a descriptive partial least squares (PLS) correlation was performed using the pyls library in Python (https://github.com/rmarkello/pyls). PLS is a multivariate statistical method that identifies associations between two variable sets by extracting latent variables that maximize shared variance (Mihalik et al., 2022). We investigated how MAIA-2, STAI Y-B, and BFI-N scores were associated with the second-degree polynomial coefficients (quadratic, linear and intercept) modeling alpha-frequency. All data were transformed using the Yeo-Johnson power function and standardized before PLS regression. Given the 10 psychological variables (8 MAIA-2 dimensions, plus STAI Y-B and BFI-N) and 192 coupling coefficients (3 polynomial coefficients × 64 electrodes), 10 sets of weights and 10 pairs of latent variables were extracted. Significance of latent variable associations was assessed via 5000 permutations; component weight stability was evaluated using the 95% confidence interval (CI) estimated by 5000 bootstraps. This analysis was run, independently with HA and IBS. Then, each of the corresponding extracted components were compared between groups.

To robustly compare gastric-brain coupling correlations with psychophysiological indices between IBS and HA, bootstrap estimates of Spearman correlations (10 000 resamples) were computed within each group. Group differences were obtained from the bootstrapped distribution of correlation differences. The two-tailed p-value was calculated as: *p* = 2 × min(perc, 1-perc), where perc is the percentile rank of zero in the bootstrapped distribution.

### 2.7. Control Analyses

Because phase-amplitude coupling quantified with the MI increases both with EEG power and the synchronization of EEG power bursts with gastric slow wave phase, we tested whether the observed changes in gastric-brain coupling dynamics across modules could be explained by changes in EEG power alone. This analysis aimed to isolate coupling effects specifically attributable to gastric-brain rhythmic synchronization. To do so, within-subject mediation analyses were conducted using the JSMediation package in R (Batailler et al., 2019). Gastric-brain coupling strength was log-transformed (real/chance ratio) to meet normality and linearity assumptions. Mediation significance was evaluated per electrode and corrected for multiple comparisons using FDR correction for each module comparisons (see Supplementary Materials for detailed explanations). Moreover, partial Spearman correlations between gastric-brain coupling and physiological variables with EEG power as a covariate were assessed to evaluate the independence to EEG power of the gastric-brain correlation with physiological variables. This was done with the Pingouin Python package (Vallat, 2018).

## 3. Results

### 3.1. Gastric-Brain Coupling

#### 3.1.1. Gastric-Brain Coupling Across Cortical Rhythms

We first checked if gastric-brain coupling existed beyond alpha rhythm. Gastric-brain coupling involved frequency-electrode pairs ranging from 1 to 45 Hz and broadly distributed across the scalp for HA (*t_mean_* = 2.70, *p* < .0001, N*_electrodes_* = 64). We then specifically tested the existence of the gastric-brain coupling strength over classical cortical rhythms for HA and IBS. Gastric-brain coupling was significantly higher than chance in delta 1-4 Hz, theta 4-8 Hz, alpha 8-13 Hz, beta 13-30 Hz and low gamma 30-45 Hz (all clusters *p* < .0001, see Supplementary Table 1 for detailed cluster statistics) with a different spatial repartition of the gastric-brain coupling per frequency band (Figure 2). No significant difference was observed between HA and IBS for each cortical rhythm (Supplementary Table 1).

**Figure 2.**
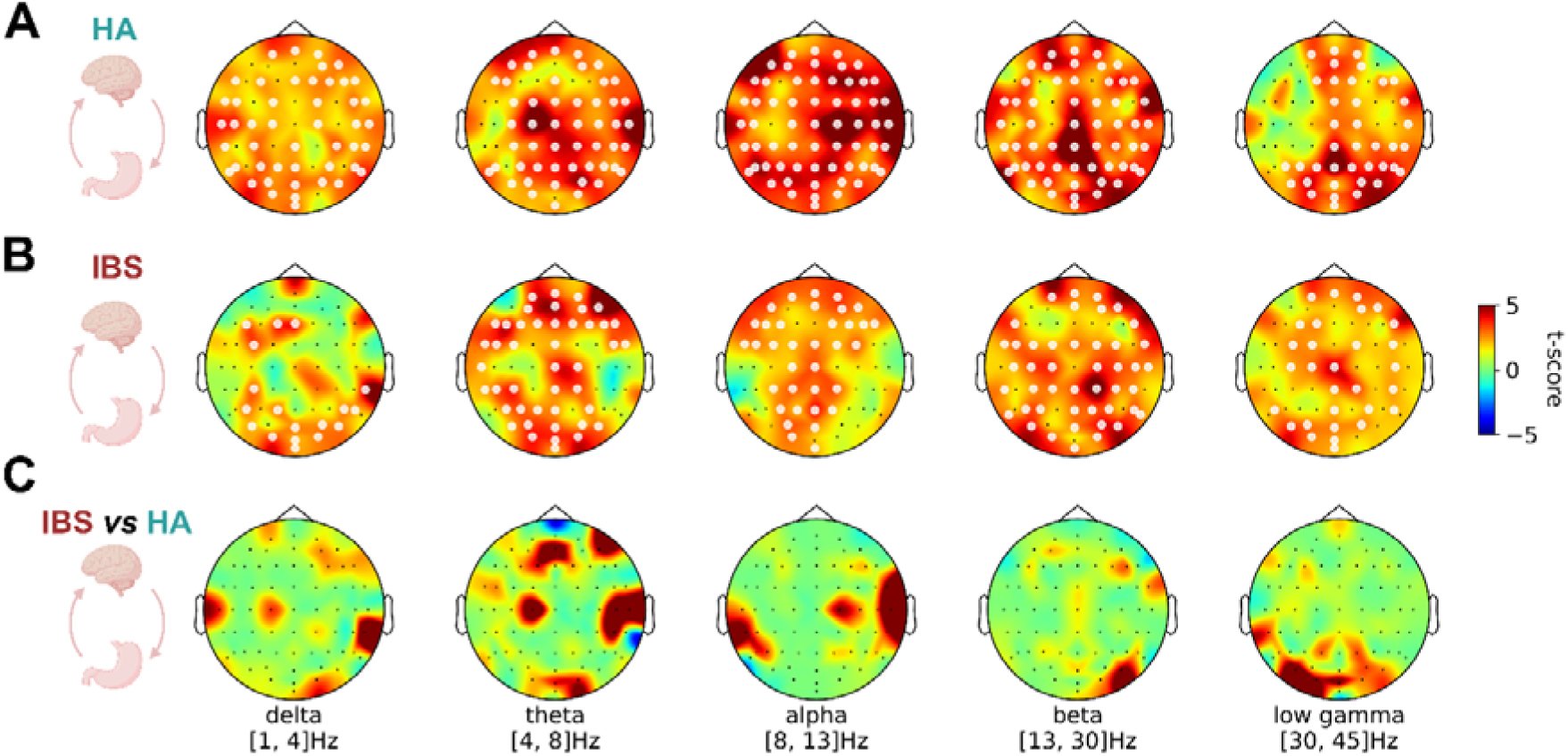
Topographic representation of the spatial cluster of gastric-brain coupling against chance level coupling per cortical rhythms for the whole session 1 and 2. A, for Healthy Adults (HA); B, for Irritable Bowel Syndrome (IBS); C, for IBS versus HA. White dots highlight electrodes included in a significant cluster (p_cluster_ < .05). Pictograms were drawn from BioRender.

#### 3.1.2. Gastric-brain Coupling Dynamics in Reactivity and Recovery

Among all the cortical rhythms coupled to gastric slow waves, the gastric-brain coupling in HA, was most sensitive to task-related reactivity and recovery in the 10 and 13 Hz alpha rhythm over left temporal, central, and right occipito-parietal electrodes (*F_mean_* = 5.04, *F_min_* = 2.46, *F_max_* = 11.16, *p* < .0001, N*_electrodes_* = 61; Figure 3A). No significant main effect of session or module-by-session interaction was observed as no significant cluster was identified. This suggests a similar effect of the module on gastric-brain coupling across the two sessions. Post-hoc analyses were then performed on gastric-brain coupling averaged in 10-13 Hz. In both HA and IBS coupling significantly decreased during the task across the scalp, with weaker effects over bilateral central electrodes, increased during all three recovery segments (see detailed post-hoc statistics in Supplementary Table 2). Thus, alpha gastric-brain coupling all over the scalp, followed a quadratic temporal trajectory in both groups (Figure 3B). With the quadratic fit *f* as a function of time *t* in minutes of the whole scalp gastric-brain coupling averaged over the 10-13 Hz and t = 0 min corresponding to the mid-point of the resting module (middle time of [10, 20] min), the fitted trajectories were : *f*_HA_(*t*) = 2.1×10^-6^ × *t*² - 1.8×10^-4^ × *t* + 3.2×10^-3^ and *f*_IBS_(*t*) = 2.4×10^-6^ × *t*² - 1.8×10^-4^ × *t* + 2.5×10^-3^. Concerning the spatial dynamics, gastric-brain coupling remained lower than baseline during the first 10 min of recovery in bilateral centro-parietal, occipital, and right frontal electrodes before returning to baseline state levels after 10 min of recovery (Figure 3C). No significant difference in gastric-brain coupling strength was observed between HA and IBS in any module (all *p_cluster_* > .05; Supplementary Table 2).

**Figure 3.**
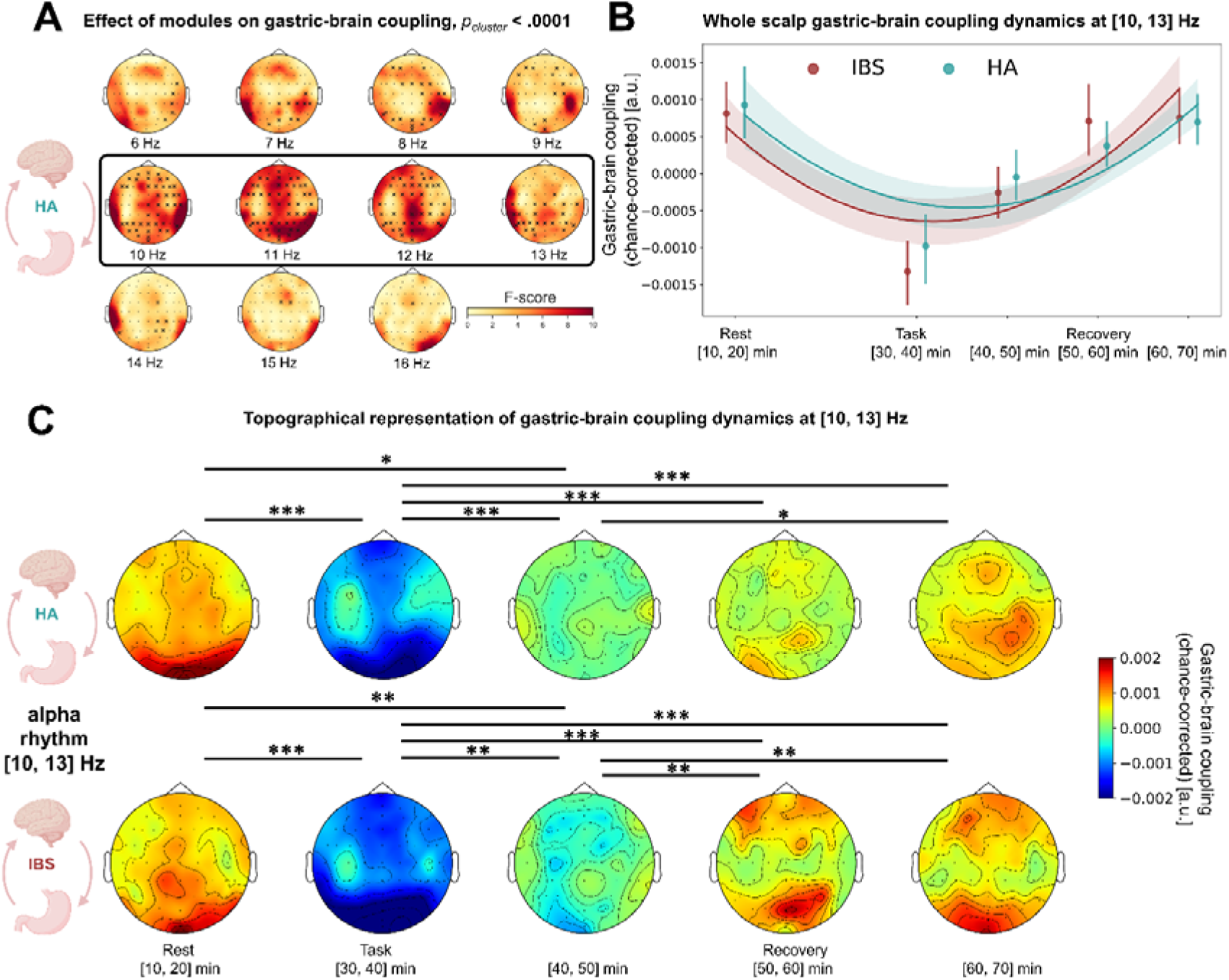
Temporal dynamics of gastric-brain coupling in Healthy Adults (HA) and Irritable Bowel Syndrome (IBS). A, Topographic representation of the significant cluster of the effect of modules on gastric-brain coupling. Black crosses highlight the EEG channels included in the cluster and the black squared topographic maps highlight the main frequencies involved in the cluster. B, Superimposition of the quadratic fit of the whole scalp grand-averaged gastric-brain coupling at 10-13 Hz (continuous line ± 95% CI) with the corresponding chance-corrected gastric-brain coupling per module (mean ± 95% CI). C, Topographic representation of the chance-corrected gastric-brain coupling averaged at 10-13 Hz per module and their pairwise significant differences for HA and IBS. No significant difference was observed on gastric-brain coupling dynamics between IBS and HA. *: p_cluster_ < .05; ** : p_cluster_ < .01; *** : p_cluster_ < .001, Holm-Bonferroni corrected for multiple comparison. Pictograms were drawn from BioRender.

#### 3.1.3. Control Analyses on Gastric-Brain Coupling Dynamics Effects

As the phase-amplitude coupling depends both on gastric slow waves and EEG power changes, we checked if these differences could be simply explained by changes among these variables. However, it is not likely to be the case for gastric slow-waves dynamics which remained stable across modules. Indeed, no significant effect of module was observed in both groups for gastric slow-waves power (module effect: χ*²(*4) = 4.76, *p* = .31, η*_p_²* = .007; module × group effect: χ*²*(4) = 3.17, *p* = .53, η*_p_²* = .005), gastric slow-waves frequency (module effect: χ*²*(4) = 1.37, *p* = .85, η*_p_²* = .002; module × group effect: χ*²*(4) = 3.20, *p* = .53, η*_p_²* = .005), nor for slow-waves cycle duration variability (module effect: χ*²*(4) = 7.37, *p* = .12, η*_p_²* = .012; module × group effect: χ*²*(4) = 8.93, *p* = .06, η*_p_²* = .014; see Supplementary Table 4). These results highlight the stability of the gastric pace-setter slow-waves dynamics across modules.

Concerning the effect of EEG power at 10-13 Hz on gastric-brain coupling dynamics, we found for both HA and IBS, that gastric-brain dynamics remained significant in occipital, temporal and central electrodes after controlling for EEG power dynamics. This suggests that these regions were mainly driven by gastric-brain coupling time-synchronization. However, we observed a distinct pattern for HA and IBS in frontal and parietal regions. In HA, EEG power partly explained the recovery of gastric-brain coupling in fronto-parietal regions while in IBS, EEG power fully explained the decrease in gastric-brain coupling from rest to task over fronto-central and parietal electrodes but not the subsequent recovery. Overall, gastric-brain coupling dynamics mainly reflected the synchronization of alpha EEG rhythm with gastric slow-waves, although transient changes in alpha power partly explained the task-to-recovery dynamics in HA and the rest-to-task dynamics in IBS. Detailed statistics and figures for this analysis are further detailed in the Supplementary Table 3 and Supplementary Figure 1.

### 3.2. Gastric-Brain Coupling Autonomic Correlates

#### 3.2.1. Autonomic Dynamics in Reactivity and Recovery

The effect of modules on EDA-SCR was significant (χ*²*(4) = 206.8, *p* < .001, η*_p_²* = .25). No significant effect of group (χ*²*(1) = 0.32, *p* = .57, η*_p_²* = .005) nor interaction effect between modules and group (χ*²*(4) = 3.93, *p* = .42, η*_p_²* = .25) were observed. Post-hoc pairwise comparisons revealed that EDA-SCR amplitude significantly increased from rest to task, remained elevated during the first 10 min of recovery and then gradually decreased toward baseline similarly in IBS and HA (Figure 4A, see Supplementary Table 4-5).

**Figure 4.**
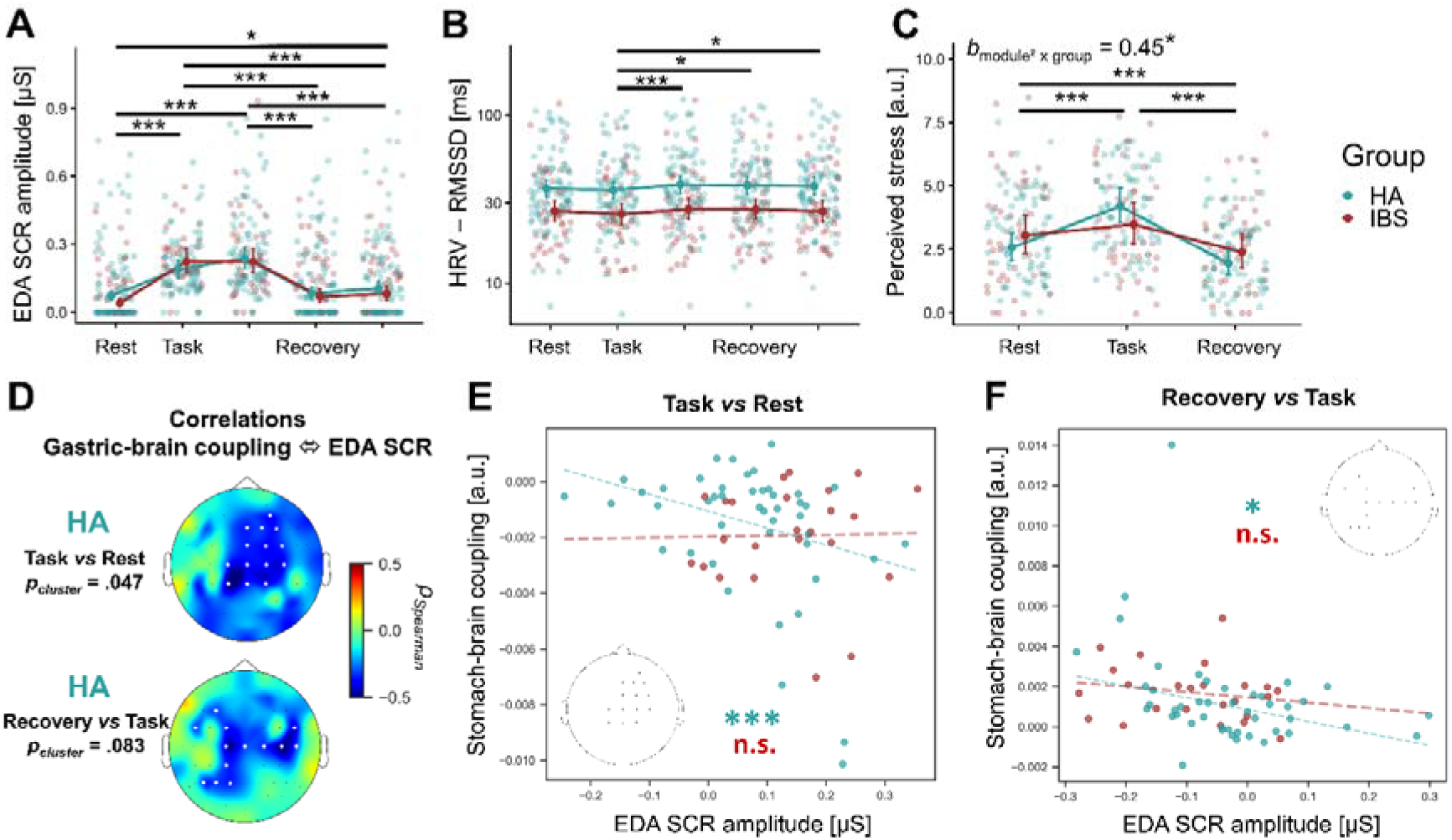
A, EDA-SCR dynamics, B, HRV-RMSSD displayed on a log-transformed y-axis, C, Perceived stress for Healthy Adults (HA) and Irritable Bowel Syndrome (IBS). The estimated marginal mean and the 95% CI are superimposed to individual values for the two experimental sessions. D. Topographic map of the two strongest clusters of Spearman correlation between gastric-brain coupling and EDA-SCR dynamics. White dots highlight the electrodes included in the cluster. E, F. Scatter plot and robust linear fit of the gastric-brain coupling changes averaged on the corresponding electrodes displayed on the head model per group. The significance of the association is represented per group (red for IBS, blue for HA). n.s.: non-significant, *: p < .05; ** : p < .01; *** : p < .001. Pictograms were drawn from BioRender.

The effect of modules on HRV-RMSSD was significant (χ*²*(4) = 17.29, *p* = 0.002, η*_p_²* = .027). IBS had an overall lower HRV-RMSSD than HA (χ*²*(1) = 5.98, *p* = 0.014, η*_p_²* = .080). However, there was no significant interaction between modules and groups (χ*²*(4) = 4.11, *p* = 0.39, η*_p_²* = .007). Post-hoc pairwise module comparisons showed that HRV-RMSSD was significantly higher during all three recovery periods relative to the task (recovery 0-10 min: *t*(639.5) = 3.63, *p_Tukey_* = .003; recovery 10-20 min: *t*(639.5) = 3.12, *p_Tukey_* = .016; recovery 20-30 min: *t*(639.5) = 3.13, *p_Tukey_* = .015). No significant difference was observed in task or recovery periods compared to rest (all *p_Tukey_* > .20, Figure 4B). Overall, HRV-RMSSD increased from task to recovery similarly in IBS and HA (see Supplementary Table 4-5 for detailed statistics).

#### 3.2.2. Association between Gastric-Brain Coupling and Autonomic Dynamics

Gastric-brain coupling dynamics were significantly negatively correlated to EDA-SCR amplitude changes from rest to task over right-lateralized fronto-central electrodes for HA (cluster ρ*_mean_* = -0.37, *p* = .047, N*_electrodes_* = 14, Figure 4D). Similarly, gastric-brain coupling dynamics revealed a trend toward a negative correlation with changes in EDA-SCR amplitude over bilateral centro-parietal electrodes from the task to the 20-30 min recovery (cluster ρ*_mean_* = -0.33, *p* = .083, N*_electrodes_* = 13, Figure 4D). These results highlight an opposite relationship between gastric-brain coupling and EDA-SCR dynamics.

The distinction between HA and IBS on the association between gastric-brain coupling and EDA-SCR dynamics was further tested by averaging gastric-brain coupling on the two clusters of electrodes identified (see Figure 4D) and by computing the correlations with EDA-SCR between the two groups. The gastric-brain coupling correlation with EDA-SCR from rest-to-task was significantly more negative for HA than for IBS (ρ*_IBS_* = 0.15, *p_bootstrap_* = .51; ρ*_HA_* = -0.41, *p_bootstrap_* = .005; ρ*_IBS-HA_* = 0.56, *p_bootstrap_* = .032, Figure 4E). Similarly, the gastric-brain coupling correlation with EDA-SCR from task-to-recovery during the final 10 min of recovery, was descriptively more negative for HA than for IBS although the between-group difference did not reach statistical significance (ρ*_IBS_* = -0.16, *p_bootstrap_*= .49; ρ*_HA_* = -0.40, *p_bootstrap_* = .013; ρ*_IBS-HA_* = 0.238, *p_bootstrap_*= .305; Figure 4F).

Despite the concomitant increase in HRV-RMSSD and gastric-brain coupling during recovery, these two variables were not significantly correlated in HA and IBS (all clusters *p* > .10).

#### 3.2.3. Control Analyses of Gastric-Brain Association with Electrodermal Activity

These associations could not be simply explained by EEG power changes. Indeed, gastric-brain coupling negative correlations with EDA-SCR were significant even when controlling for EEG power changes for healthy adults (task *vs* rest: ρ*_HA_* (43) = -0.43, *p* = .004; recovery *vs* task: ρ*_HA_* (43) = -0.38, *p* = .011). Additionally, for IBS, the strength of association was similar or indicated an even lower association than HA after adding EEG power as covariate (task *vs* rest: ρ*_IBS_* (22) = 0.11, *p* = .61; recovery *vs* task: ρ*_IBS_* (22) = -0.16, *p* = .47). Thus, the negative correlations between gastric-brain coupling and EDA-SCR and their different strength in IBS and HA are independent of EEG power changes.

### 3.3. Gastric-Brain Coupling and Psychological Variables

#### 3.3.1. Association Between Gastric-Brain Coupling and Perceived Stress

The effect of module on perceived stress was significant (χ*²(*2) = 56.77, *p* < .001, η*_p_²* = .175) as well as the interaction between modules and group (χ*²(*2) = 6.97, *p* = .031, η*_p_²* = .022) with no significant overall difference between groups (χ*²(*1) = 1.05, *p* = .30, η*_p_²* < .001). Post-hoc paired contrasts on module effect showed that perceived stress increases from rest to task (*t*(325.4) = 4.10, *p_Tukey_* < .001) and decreases from task to the end of recovery (*t*(325.4) = - 8.06, *p_Tukey_* < .001) and reached to a lower level than rest (*t*(325.4) = 3.96, *p_Tukey_* < .001). Post-hoc polynomial contrasts on module × group interaction showed that perceived stress variations across modules was significantly attenuated in IBS compared to HA (quadratic coefficient for IBS *vs* HA: *t*(325.4) = 2.52, *p_Holm_* = .024, Figure 4C, see Supplementary Table 4-5 for detailed statistics). No significant correlation between gastric-brain coupling dynamics and perceived stress were observed (all clusters *p* > .10).

#### 3.3.2. Association Between Gastric-Brain Coupling, Interoception, Anxiety and Neuroticism

The descriptive PLS-correlation analysis conducted in HA revealed a significant association between a latent psychological variable (i.e., constructed from anxiety, neuroticism, and interoceptive awareness) and a latent gastric-brain coupling quadratic dynamics variables (*r* = .52, *p* = .004, 5000 permutations across 10 components; Figure 5A). Anxiety and neuroticism traits positively and reliably load to the psychological latent variable while “Body-listening”, “Emotional Awareness”, “Self-Regulation” and “Noticing” dimensions of interoceptive awareness (MAIA-2) negatively and reliably load to this psychological latent variable (Figure 5B). In other words, higher scores on the psychological latent variable indicate a tendency to experience greater anxiety and negative emotions without listening and using interoceptive feelings to regulate emotions. EEG electrodes F1, C1, CP1, CPz, and CP4 positively and stably loaded on the gastric-brain coupling dynamic latent variable for the quadratic coefficient and CP1 and CP4 negatively and stably loaded on the gastric-brain coupling dynamic latent variable for the linear coefficient (Figure 5C). Higher scores on the gastric-brain coupling dynamics latent variable indicated higher gastric-brain coupling reduction from rest to task and higher gastric-brain coupling augmentation from task to recovery. As these two latent variables positively correlated for HA, higher anxiety and neuroticism traits with lower tendency to use interoceptive feeling to regulate one’s emotion were associated to stronger gastric-brain centro-parietal dynamical variations in response to and recovery from the task (Figure 5D). When extracting these exact components in IBS, a significant negative correlation between the psychological dimensions and gastric-brain coupling dynamics was observed (ρ*_IBS_* = -0.57, *p_bootstrap_* = .005) that is significantly different from that of HA (ρ*_IBS-HA_* = -1.04, *p* < .001; see Figure 5E).

**Figure 5:**
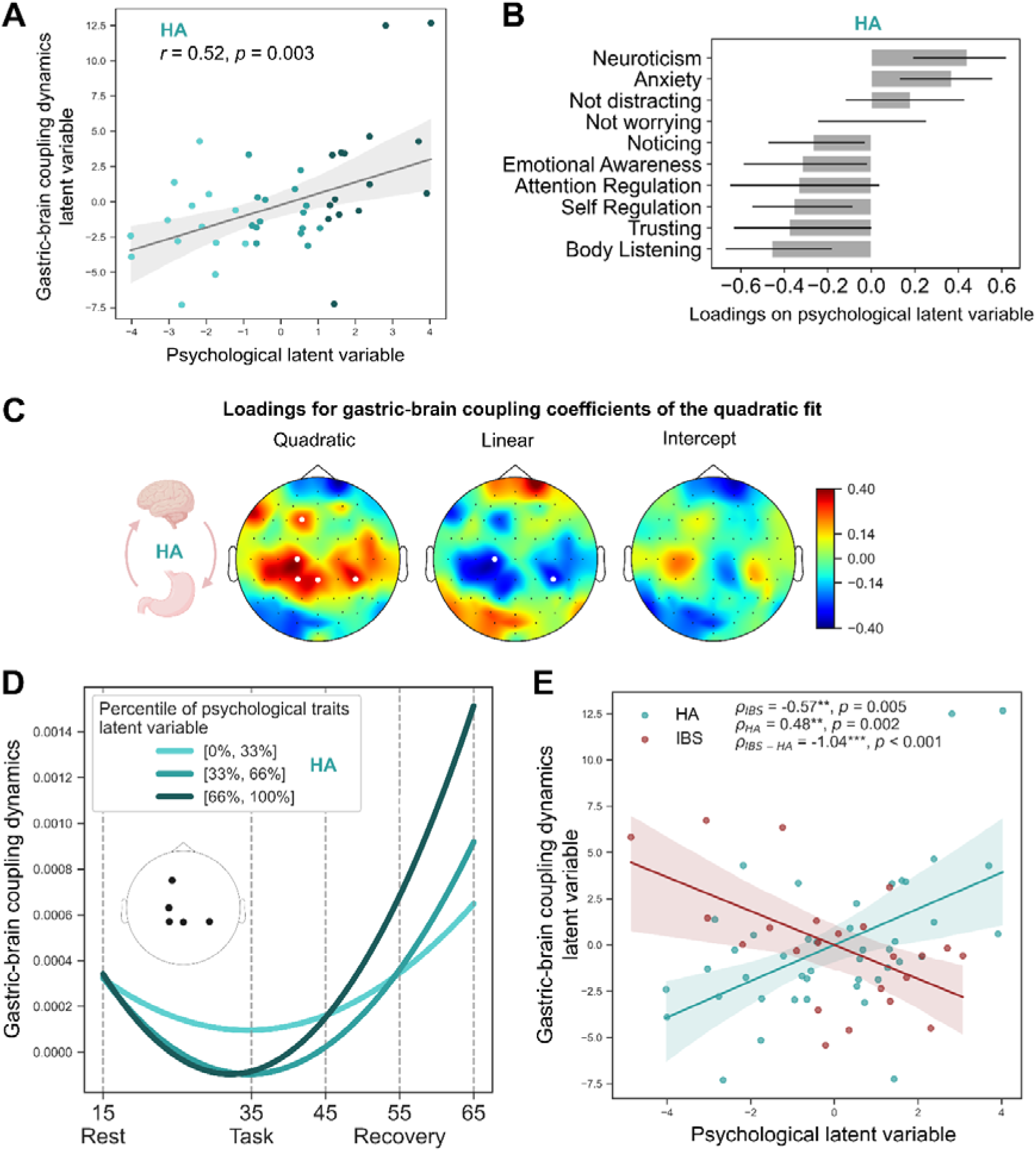
Descriptive PLS correlation model fitted in Healthy Adults (HA) and applied to Irritable Bowel Syndrome (IBS) participants. A. Scatter plot and linear regression fit showing the association between the psychological latent variable and the gastric-brain coupling dynamics latent variable in HA participants. The blue color gradient indicates participants belonging to the terciles of the psychological latent variable. B. Loadings and 95% confidence intervals (CI) of the psychological dimensions on the psychological latent variable. C. Loadings and 95% CI of the coefficients from the quadratic model describing gastric-brain coupling dynamics across modules on the gastric-brain coupling dynamics latent variable. D. Mean gastric-brain coupling dynamics averaged across the electrodes indicated by black dots, shown separately for the terciles of the psychological latent variable in HA participants. E. Scatter plots and linear regression fits (95% CI) illustrating the relationship between the psychological and gastric-brain coupling dynamics latent variables in HA and IBS participants, together with the comparison of their corresponding Spearman correlation coefficients. Pictograms were drawn from BioRender.

We next applied PLS correlation analysis in IBS participants to identify associations between psychological variables and gastric-brain coupling dynamics that may be specific to IBS. The corresponding descriptive PLS-correlation analysis revealed a significant association between a latent psychological variable and a latent gastric-brain coupling dynamics variables (*r* = .88, *p* = .017, 5000 permutations across 10 components; Figure 6A). The interoceptive awareness dimensions “Noticing”, “Emotional awareness”, “Body listening” and “Not-distracting” as well as anxiety trait and neuroticism negatively and reliably loaded on the psychological latent variable (Figure 6B). In other words, higher scores on the psychological latent variable indicate a tendency to experience lower anxiety and negative emotions with a low interoceptive awareness. EEG electrodes F7 and P9 negatively and stably loaded on the gastric-brain coupling dynamic latent variable for the intercept (i.e., rest module; Figure 6C). In other words, higher scores on the gastric-brain coupling dynamics latent variable indicate lower gastric-brain coupling at rest (Figure 6D). As these two latent variables are positively correlated only in IBS, higher interoceptive awareness with anxiety and neuroticism tendencies is associated with stronger gastric-brain coupling in frontal and parietal left lateralized electrodes. When extracting these exact components in HA, no significant correlation between these psychological variables and gastric-brain coupling at rest was observed (ρ*_HA_* = -0.04, *p_bootstrap_* = .801) and significantly differed from that of HA (ρ*_IBS-HA_* = 0.90, *p_bootstrap_*< .001; see Figure 6E).

**Figure 6:**
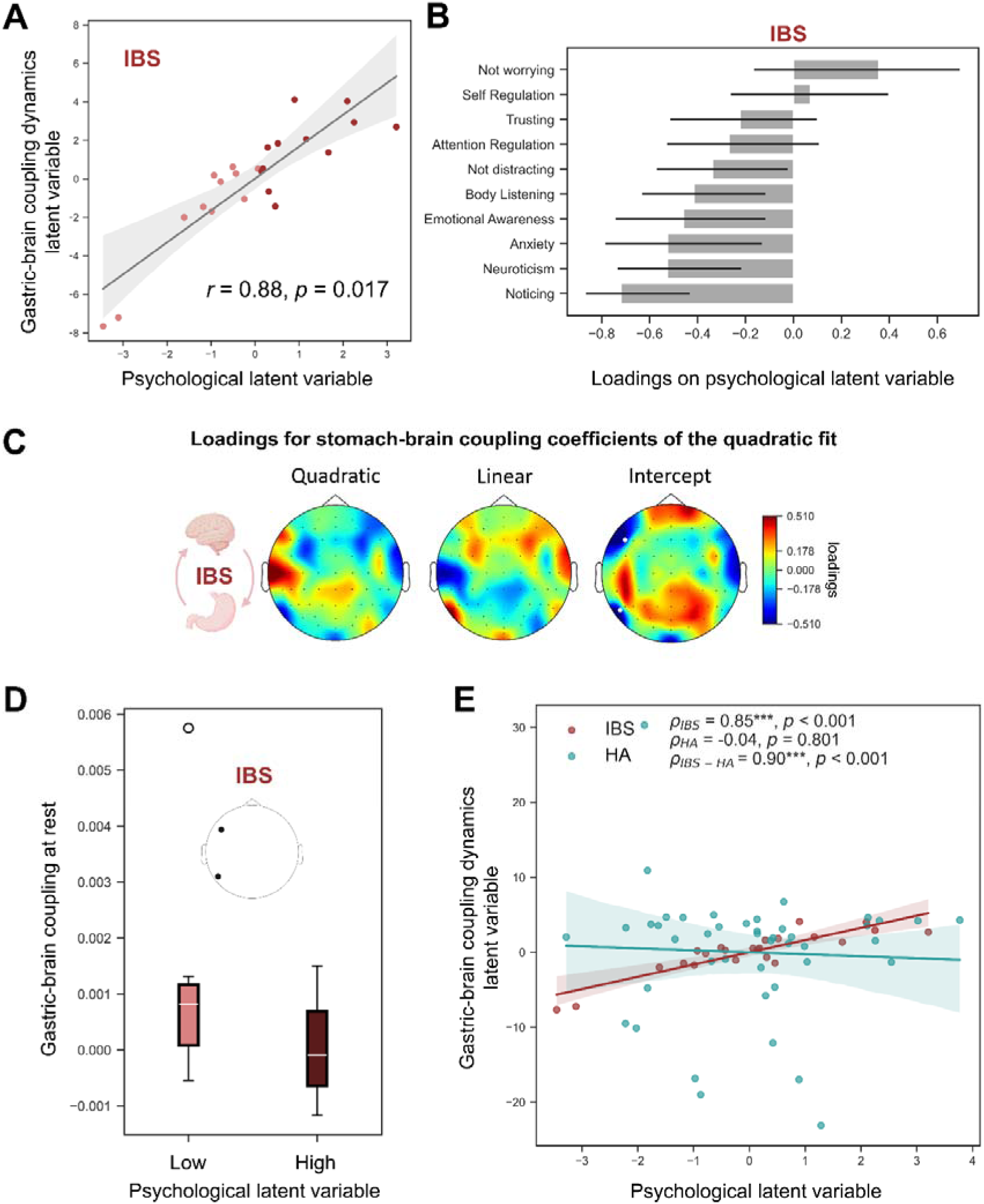
Descriptive PLS correlation model fitted in Irritable Bowel Syndrome (IBS) participants and applied to Healthy Adults (HA). A. Scatter plot and linear regression fit showing the association between the psychological latent variable and the gastric-brain coupling dynamics latent variable in IBS participants. The red color gradient indicates participants belonging to the low- and high-median-split groups defined according to the psychological latent variable. B. Loadings and 95% confidence intervals (CI) of the psychological dimensions on the psychological latent variable. C. Loadings and 95% CI of the coefficients from the quadratic model describing gastric-brain coupling dynamics across modules on the gastric-brain coupling dynamics latent variable. D. Mean gastric-brain coupling dynamics averaged across the electrodes indicated by black dots, shown separately for the low- and high-median-split groups on the psychological latent variable in IBS participants. E. Scatter plots and linear regression fits (95% CI) illustrating the relationship between the psychological variables and gastric-brain coupling dynamics latent variables in HA and IBS participants, together with the comparison of their corresponding Spearman correlation coefficients. Pictograms were drawn from BioRender.

#### 3.3.3. Control Analyses on Gastric-Brain Coupling Dynamics Psychological Correlates

The PLS-correlation findings applied on HA are unlikely to be explained by EEG power dynamic changes as no significant associations were found in HA between psychological variables and EEG power dynamics when applying the same PLS correlation methodology (all *p* > .33). Additionally, when applying the PLS-correlation weights obtained for the gastric-brain coupling dynamics on EEG coefficients for IBS, no significant association between psychological variables and EEG power dynamics were observed (ρ*_IBS_* = -0.37, *p_bootstrap_* = .083). In addition, no significant difference between groups was observed on the association between psychological variables and EEG power dynamics (ρ*_IBS-HA_* = -0.40, *p_bootstrap_* = .12). Taken together, these analyses confirm that the distinct relationship between gastric-brain coupling and psychological variables for HA and IBS reflects a neurovisceral integration profile based on gastric-brain rhythms synchrony.

The PLS-correlation findings applied on IBS are unlikely to be explained by EEG power dynamics alone, as no significant associations were observed in IBS between psychological variables and EEG power dynamics using the same PLS correlation approach (all *p* > .12). In addition, the psychological latent variable was primarily associated with gastric-brain coupling at rest. As gastric-brain coupling may also be influenced by gastric slow-waves, we therefore tested whether this relationship was independent of gastric slow-wave characteristics, including power, cycle duration, and variability of cycle duration. In both IBS and HA groups, the association between latent variables remained stable after controlling for these gastric measures (ρ*_IBS_* = 0.83 to 0.85, all *p* < .0001; ρ*_HA_* = −0.04 to −0.025, all *p* > .79).

All distinct relationships for psychological variables with gastric-brain coupling observed between IBS and HA could not be explained by significant differences on psychological variables between IBS and HA (see Supplementary Table 6).

Taken together, these control analyses suggest that IBS status specifically moderates the relationship between gastric-brain coupling and anxiety, neuroticism and interoceptive awareness. Indeed, these relationships cannot be simply explained by psychological, EEG power or gastric slow waves dynamics differences between HA and IBS.

## 4. Discussion

The present study investigated the functional role of phase-amplitude gastric-brain coupling by examining its psychophysiological correlates in HA and adults with a disorder of gut-brain interactions, namely IBS, at rest and under cognitive effort. We first showed that gastric-brain coupling spanned a broad range of cortical rhythms and scalp regions, but gastric-alpha cortical rhythm coupling showed the strongest contextual modulation, decreasing during a cognitive-emotional challenge and progressively recovering afterward. Second, decrease in gastric-brain coupling was primarily associated with sympathetic activation rather than parasympathetic withdrawal. Third, although average gastric-brain coupling dynamics did not differ between IBS and HA, interoception, anxiety and neuroticism as well as autonomic correlates of coupling differed between these groups. Together, these findings support a functional role of gastric-brain coupling in the integration of interoceptive, autonomic, and affective processes and further refine current models of gut-brain-mind interactions.

### 4.1. Functional Role of Gastric-Brain Coupling Dynamics

Across conditions, the dynamic changes in gastric-brain coupling strength were driven by phase synchronization between gastric slow waves and cortical activity rather than by changes in gastric rhythm characteristics or cortical alpha power. This suggests that gastric-brain coupling is an actively regulated coordination mechanism rather than a passive consequence of fluctuations in either gastric or cortical activity. During a demanding cognitive task, the brain transiently disengages its temporal alignment with the stomach’s intrinsic rhythm, allowing flexible adaptation to externally driven processing demands.

Among frequency bands, gastric-cortical alpha coupling was the most sensitive to contextual change, indicating a privileged role for alpha rhythm in gastric-cortical alignment. Alpha rhythms are increasingly recognized as a fundamental mechanism for regulating cortical excitability and information flow. A recent synthesis further supports alpha as a mechanistic substrate for rhythmic inhibition, controlling excitability, and implementing periodic sampling of sensory input across cortical regions (Pascucci et al., 2025). From this perspective, gastric-alpha coupling may reflect a dynamic alignment between a dominant interoceptive bodily oscillator and cortical inhibitory sampling mechanisms, potentially enabling periodic access to internal bodily states. Our results suggest that such alignment characterizes resting-state physiology, whereas cognitive engagement temporarily interrupts this synchronization, presumably to prioritize processing of externally relevant information. Importantly, this reduction in coupling was not accompanied by changes in gastric slow waves, indicating that coupling reflects temporal coordination with cortical alpha bursts rather than changes in the gastric pacemaker. This observation strengthens the view that gastric slow waves provide a reliable physiological reference signal upon which cortical dynamics can flexibly synchronize or desynchronize depending on behavioural context (Engelen et al., 2023).

Additionally, the recovery of gastric-brain coupling toward baseline levels is aligned with the emerging hypothesis that gastric-brain coupling may support the organism’s energy consumption and storage strategies (Rebollo et al., 2021). For example, gastric-brain coupling intensity has been shown to positively correlate with weight loss (Levakov et al., 2021). In line with theories of interoceptive inference (Seth, 2013), stronger gastric-brain coupling may help reduce energetic costs by increasing the predictability of cortical activity and lowering cortical entropy (Bettinger & Friston, 2023), as cortical activity becomes temporally aligned with a stable and predictable visceral oscillator. Interrupting this synchronized state during task performance may reflect the energetic demands of task engagement, whereas rest and recovery may correspond to energetically efficient viscera-brain synchronized states.

### 4.2. Autonomic Modulation of Gastric-Brain Coupling

Bidirectional gut-brain interactions are mediated by the ANS, primarily via vagal and spinal pathways (Lane et al., 2009). While prior work has shown that resting gastric-brain coupling relates to combined autonomic influences (Rebollo & Tallon-Baudry, 2022) and can be modulated by vagal stimulation (Müller et al., 2022), its contextual modulation across cognitive-emotional states remained unclear. Here, coupling changes were primarily associated with sympathetic activation, as indexed by EDA, whereas parasympathetic activity, as indexed by RMSSD, was not associated with coupling dynamics. Specifically, increased sympathetic arousal was associated with reduced gastric-alpha rhythm coupling, particularly in centro-parietal regions. These effects are anatomically consistent with recent findings in rodents highlighting somatosensory and motor cortices on sympathetic influences on the stomach (Levinthal & Strick, 2020). Although interpreting this result as a sympathetic top-down inhibition of gastric function is plausible, the absence of direct changes in gastric slow-wave amplitude or frequency limits mechanistic inference at the peripheral level. Future studies using gastric pills to quantify gastric acid secretion (Porciello et al., 2024) across contextual changes, as well as direct measures of gastric muscular contractions, may help disentangle this hypothesis.

Given that alpha rhythms reflect thalamo-cortical loops (Halgren et al., 2019), that the thalamus integrates visceral signals and relays them to cortical regions (Rebollo et al., 2021), and that EDA-SCR indices are shaped by cortical and subcortical processes involved in cognitive and emotion (Sequeira et al., 2009), our findings may be consistent with top-down corticothalamic inhibition of the thalamic relay of gastric inputs to the cerebral cortex during physiological arousal. From this perspective, the gastric rhythm appears more as a source of perturbation than as a facilitator of cortical activity under cognitive demand.

### 4.3. Gastric-Alpha Coupling, Interoceptive Awareness, and Negative Affectivity

We explored whether the dynamical evolution of gastric-brain coupling through contextual changes more than its resting-state level could be a potential physiological mechanism indexing individual differences in emotional and interoceptive awareness (Engelen et al., 2023; Fermin et al., 2022; Seth, 2013). We observed that maintaining gastric-brain coupling at levels closer to those observed during rest was associated with higher interoceptive abilities, lower anxiety and a reduced tendency to feel negative emotions. This may reflect a functional mechanism of gastric-brain coupling during the task, that enables periodic sampling of internal bodily signals, supporting adaptive emotional regulation during cognitive demands (Fermin et al., 2024; Seth, 2013). Conversely, reduced coupling may index diminished access to interoceptive signals under high arousal.

### 4.4. Altered Psychophysiological Organization in Irritable Bowel Syndrome

Although gastric-brain coupling dynamics was comparable between IBS and HA, the functional embedding of this coupling within autonomic and psychological dimensions differed substantially. On the autonomic side, in IBS, a stronger sympathetic arousal during the task was not related to a stronger reduction in gastric-brain coupling, contrary to what was observed in HA. This may reflect a persistent influence of gastric rhythm over cortical activity in IBS, even during heightened arousal, that can be seen as a dysfunctional top-down regulation of gastric inputs in the context of emotional and cognitive demands. On the psychological side, gastric-brain coupling dynamics were oppositely related to psychological variables in IBS compared to HA. These exploratory results may index a particular gastric-brain-mind integration in IBS, shaping how interoceptive signals are integrated during emotional and cognitive challenges. Moreover, we observed at rest, that IBS participants showed stronger associations than HA, between coupling and anxiety/interoceptive awareness, particularly in left-lateralized frontal and parietal regions, suggesting altered cortical representation of visceral signals. Recently, resting state gastric-brain coupling was shown, in a large cohort, to increase with a composite profile primarily characterized by anxiety, depressive symptoms, and poor mental well-being, and, to a lesser extent, by interoceptive awareness (Banellis et al., 2025). We extend these results in a model of gut-brain axis disorder by showing that resting-state gastric-brain coupling is more strongly associated with interoceptive awareness than in HA. This finding aligns with models of functional gastrointestinal disorders that emphasize disrupted interoceptive processing, alexithymia-like traits, and altered neurovisceral integration (De Gucht & Heiser, 2003; Kano et al., 2018). It further suggests that gastric-brain coupling may provide a stronger index of this disruption in IBS than in healthy individuals. More broadly, our exploratory findings encourage future research to examine not only how static and resting-state gastric-brain coupling relate to psychological variables, but also how the dynamical trajectories of brain-body states relate to psychology are associated with psychological functioning in disorders of gut-brain axis (Kluger et al., 2024).

### 4.5. Limitations and Conclusion

Although our task reliably induced arousal, more ecologically valid paradigms may reveal additional coupling dynamics relevant in everyday challenges. The absence of correlation with a parasympathetic cardiac index and perceived stress may limit the understanding of the neurovisceral regulation function of gastric-brain coupling. Finally, despite the broad spatial and spectral distribution of gastric-brain coupling, psychophysiological correlates were observed primarily within alpha rhythms and central scalp regions, suggesting that gastric-brain synchrony may serve multiple functional roles depending on its spatial and spectral characteristic, which remain to be elucidated. Moreover, the exploratory findings on the psychological association with gastric-brain coupling dynamics need to be replicated in a larger sample.

In conclusion, gastric-cortical coupling transiently decreases during physiological arousal. Its context-dependent dynamics, rather than resting-state levels alone, may reflect the integration of autonomic, interoceptive, and affective processes. Although average coupling dynamics were preserved in IBS, their altered autonomic and psychological correlates suggest disrupted gut-brain-mind integration. These findings support a functional role for gastric rhythms in coordinating cortical activity and highlight gastric-brain coupling dynamics as a promising physiological marker to explore gut-brain functioning in health and disease.

## Conflict of Interest Statement

The authors declare no conflicts of interest.

## Statement of Ethics

All participants signed an informed consent, the privacy rights of human subjects were observed, all procedures were performed in compliance with relevant laws and institutional guidelines. The study was approved by the ethics committee of University of Savoie Mont Blanc, France (reference C.E.R.E.U.S: CER_2021_10) and by the research ethics committee of SUD-EST VI Clermont-Ferrand, France (reference AU 1658: 2020-A02155-34).

## Funding Sources

This work was supported by research grants from *i)* NeuroCog IDEX UGA under the “Investing for the future program”, *ii)* FONDATION Université Savoie Mont Blanc for the “Heart-brain project” and “INGAST” project, and *iii)* ANR USMB SHINE for the “6ème Sens” project. The funder had no role in the design, data collection, data analysis, and reporting of this study.

## Author Contributions

Jeanne, R., Minjoz, S., Bonaz, B., Hot, P., Kibleur, A., and Pellissier, S., contributed to the methodology, study design, and funding acquisition. Minjoz, S., and Bonaz, B., contributed to the project administration and participant’s selection. Jeanne, R., Minjoz, S., and Sinniger, V., collected the data. Jeanne, R., and Minjoz, S., Hot, P., Kibleur, A., and Pellissier, S., performed the investigation, formal analysis and interpretation. Jeanne, R. performed the data visualization. Hot, P., Kibleur, A., and Pellissier, S., supervised the project. Jeanne, R., Minjoz, S., and Pellissier, S., drafted the manuscript. Minjoz, S., Sinniger, V., Bonaz, B., Hot, P., Kibleur, A., and Pellissier, S., provided critical revision on the manuscript. All authors reviewed and approved the final version of the manuscript.

## Data Availability Statement

The study was pre-registered on OSF (https://doi.org/10.17605/OSF.IO/FKB79) and in ClinicalTrials.gov (Identifier: NCT04807933). Once the manuscript is published, the scripts that support the findings of the study will be made openly available on the OSF project folder (https://doi.org/10.17605/OSF.IO/FKB79). Other materials and information about this study are available from the corresponding author upon reasonable request.

## Data Transparency Statement

This study is a part of a larger project that came up with two publications evaluating psychophysiological effects of heart rate variability biofeedback in irritable bowel syndrome (Minjoz et al., 2025) and healthy participants (Minjoz et al., 2026). None of the analyses and results presented in the current study have been published or considered for publication elsewhere.

## Notes

https://doi.org/10.17605/OSF.IO/FKB79

### Competing Interest Statement

The authors have declared no competing interest.

## References

Ablin, P., Cardoso, J.-F., & Gramfort, A. (2018). Faster Independent Component Analysis by Preconditioning With Hessian Approximations. IEEE Transactions on Signal Processing, 66(15), 4040–4049. 10.1109/TSP.2018.2844203

Aru, J., Aru, J., Priesemann, V., Wibral, M., Lana, L., Pipa, G., Singer, W., & Vicente, R. (2015). Untangling cross-frequency coupling in neuroscience. Current Opinion in Neurobiology, SI: Brain Rhythms and Dynamic Coordination, 31, 51–61. 10.1016/j.conb.2014.08.002

Azzalini, D., Rebollo, I., & Tallon-Baudry, C. (2019). Visceral Signals Shape Brain Dynamics and Cognition. Trends in Cognitive Sciences, 23(6), 488–509. 10.1016/j.tics.2019.03.007

Balasubramani, P. P., Walke, A., Grennan, G., Perley, A., Purpura, S., Ramanathan, D., Coleman, T. P., & Mishra, J. (2022). Simultaneous Gut-Brain Electrophysiology Shows Cognition and Satiety Specific Coupling. Sensors, 22(23), Article 23. 10.3390/s22239242

Banellis, L., Rebollo, I., Nikolova, N., & Allen, M. (2025). Gastric-brain coupling indexes a dimensional signature of mental health. Nature Mental Health, 3(8), 899–908. 10.1038/s44220-025-00468-6

Batailler, C., Muller, D., Yzerbyt, V., & Judd, C. (2019). Smediation: Mediation Analysis Using Joint Significance (p. 0.2.2) [Jeu de données]. 10.32614/CRAN.package.JSmediation

Bates, D., Mächler, M., Bolker, B., & Walker, S. (2015). Fitting linear mixed-effects models using lme4. Journal of statistical software, 67, 1–48.

Bettinger, J. S., & Friston, K. J. (2023). Conceptual foundations of physiological regulation incorporating the free energy principle and self-organized criticality. Neuroscience & Biobehavioral Reviews, 155, 105459. 10.1016/j.neubiorev.2023.105459

Boucsein, W., Fowles, D. C., Grimnes, S., Ben-Shakhar, G., roth, W. T., Dawson, M. E., Filion, D. L., & Society for Psychophysiological Research Ad Hoc Committee on Electrodermal Measures. (2012). Publication recommendations for electrodermal measurements. Psychophysiology, 49(8), 1017–1034. 10.1111/j.1469-8986.2012.01384.x

Critchley, H. D. (2002). Review: Electrodermal Responses: What Happens in the Brain. The Neuroscientist, 8(2), 132–142. 10.1177/107385840200800209

Critchley, H. D., & Harrison, N. A. (2013). Visceral Influences on Brain and Behavior. Neuron, 77(4), 624–638. 10.1016/j.neuron.2013.02.008

De Gucht, V., & Heiser, W. (2003). Alexithymia and somatisation: A quantitative review of the literature. Journal of Psychosomatic Research, 54(5), 425–434. 10.1016/S0022-3999(02)00467-1

Drossman, D. A., & Hasler, W. L. (2016). Rome IV—Functional GI Disorders: Disorders of Gut-Brain Interaction. Gastroenterology, 150(6), 1257–1261. 10.1053/j.gastro.2016.03.035

Enck, P., Aziz, Q., Barbara, G., Farmer, A. D., Fukudo, S., Mayer, E. A., Niesler, B., Quigley, E. M. M., Rajilić-Stojanović, M., Schemann, M., Schwille-Kiuntke, J., Simren, M., Zipfel, S., & Spiller, R. C. (2016). Irritable bowel syndrome. Nature Reviews Disease Primers, 2(1), 16014. 10.1038/nrdp.2016.14

Engelen, T., Solcà, M., & Tallon-Baudry, C. (2023). Interoceptive rhythms in the brain. Nature Neuroscience, 26(10), Article 10. 10.1038/s41593-023-01425-1

Fermin, A. S. R., Friston, K., & Yamawaki, S. (2022). An insula hierarchical network architecture for active interoceptive inference. Royal Society Open Science, 9(6), 220226. 10.1098/rsos.220226

Fermin, A. S. R., Sasaoka, T., Maekawa, T., Ono, K., Chan, H.-L., & Yamawaki, S. (2024). Insula-cortico-subcortical networks predict interoceptive awareness and stress resilience. Asian Journal of Psychiatry, 95, 103991. 10.1016/j.ajp.2024.103991

Gramfort, A. (2013). MEG and EEG data analysis with MNE-Python. Frontiers in Neuroscience, 7. 10.3389/fnins.2013.00267

Greco, A., Valenza, G., Lanata, A., Scilingo, E. P., & Citi, L. (2016). cvxEDA: A Convex Optimization Approach to Electrodermal Activity Processing. IEEE Transactions on Bio-Medical Engineering, 63(4), 797–804. 10.1109/TBME.2015.2474131

Halgren, M., Ulbert, I., Bastuji, H., Fabó, D., Erőss, L., Rey, M., Devinsky, O., Doyle, W. K., Mak-McCully, R., Halgren, E., Wittner, L., Chauvel, P., Heit, G., Eskandar, E., Mandell, A., & Cash, S. S. (2019). The generation and propagation of the human alpha rhythm. Proceedings of the National Academy of Sciences of the United States of America, 116(47), 23772–23782. 10.1073/pnas.1913092116

Holm, S. (1979). A simple sequentially rejective multiple test procedure. Scandinavian Journal of Statistics. Theory and Applications, 6(2), 65–70.

Holtmann, G., & Talley, N. J. (2014). The gastric-brain axis. Best Practice & Research Clinical Gastroenterology, Gastric Physiology and Pathogenesis: Evolving Concepts and Their Impact on Clinical Practice, 28(6), 967–979. 10.1016/j.bpg.2014.10.001

Jas, M., Engemann, D. A., Bekhti, Y., Raimondo, F., & Gramfort, A. (2017). Autoreject: Automated artifact rejection for MEG and EEG data. NeuroImage, 159, 417–429. 10.1016/j.neuroimage.2017.06.030

Jeanne, R., Piton, T., Minjoz, S., Bassan, N., Le Chenechal, M., Semblat, A., Hot, P., Kibleur, A., & Pellissier, S. (2023). Gut-Brain Coupling and Multilevel Physiological Response to Biofeedback Relaxation After a Stressful Task Under Virtual Reality Immersion: A Pilot Study. Applied Psychophysiology and Biofeedback, 48(1), 109–125. 10.1007/s10484-022-09566-y

Jensen, O., & Colgin, L. L. (2007). Cross-frequency coupling between neuronal oscillations. Trends in Cognitive Sciences, 11(7), 267–269. 10.1016/j.tics.2007.05.003

John, O. P., Donahue, E. M., & Kentle, R. L. (1991). Big Five Inventory (BFI). Journal of Personality and Social Psychology. https://doi.apa.org/doi/10.1037/t07550-000

Kano, M., Dupont, P., Aziz, Q., & Fukudo, S. (2018). Understanding Neurogastroenterology From Neuroimaging Perspective: A Comprehensive Review of Functional and Structural Brain Imaging in Functional Gastrointestinal Disorders. Journal of Neurogastroenterology and Motility, 24(4), 512–527. 10.5056/jnm18072

Keitel, C., Ruzzoli, M., Dugué, L., Busch, N. A., & Benwell, C. S. Y. (2022). Rhythms in cognition: The evidence revisited. The European Journal of Neuroscience, 55(11-12), 2991–3009. 10.1111/ejn.15740

Kluger, D. S., Allen, M. G., & Gross, J. (2024). Brain-body states embody complex temporal dynamics. Trends in Cognitive Sciences, 28(8), 695–698. 10.1016/j.tics.2024.05.003

Laborde, S., Mosley, E., & Mertgen, A. (2018). Vagal Tank Theory[: The Three Rs of Cardiac Vagal Control Functioning - Resting, Reactivity, and Recovery. Frontiers in Neuroscience, 12, 458. 10.3389/fnins.2018.00458

Laborde, S., Mosley, E., & Thayer, J. F. (2017). Heart Rate Variability and Cardiac Vagal Tone in Psychophysiological Research - Recommendations for Experiment Planning, Data Analysis, and Data Reporting. Frontiers in Psychology, 08. 10.3389/fpsyg.2017.00213

Lacy, B. E., Pimentel, M., Brenner, D. M., Chey, W. D., Keefer, L. A., Long, M. D., & Moshiree, B. (2021). ACG Clinical Guideline[: Management of Irritable Bowel Syndrome. Official Journal of the American College of Gastroenterology | ACG, 116(1), 17. 10.14309/ajg.0000000000001036

Lane, R. D., Waldstein, S. R., Chesney, M. A., Jennings, J. R., Lovallo, W. R., Kozel, P. J., Rose, R. M., Drossman, D. A., Schneiderman, N., Thayer, J. F., & Cameron, O. G. (2009). The rebirth of neuroscience in psychosomatic medicine, Part I: Historical context, methods, and relevant basic science. Psychosomatic Medicine, 71(2), 117–134. 10.1097/PSY.0b013e31819783be

Lenth, R. V., & Piaskowski, J. (2026). emmeans: Estimated Marginal Means, aka Least-Squares Means. https://rvlenth.github.io/emmeans/

Levakov, G., Kaplan, A., Yaskolka Meir, A., Rinott, E., Tsaban, G., Zelicha, H., Meiran, N., Shelef, I., Shai, I., & Avidan, G. (2021). Neural correlates of future weight loss reveal a possible role for brain-gastric interactions. NeuroImage, 224, 117403. 10.1016/j.neuroimage.2020.117403

Levinthal, D. J., & Strick, P. L. (2020). Multiple areas of the cerebral cortex influence the stomach. Proceedings of the National Academy of Sciences of the United States of America, 117(23), 13078–13083. 10.1073/pnas.2002737117

Lipponen, J. A., & Tarvainen, M. P. (2019). A robust algorithm for heart rate variability time series artefact correction using novel beat classification. Journal of Medical Engineering & Technology, 43(3), 173–181. 10.1080/03091902.2019.1640306

Lüdecke, D. (2018). Ggeffects Tidy Data Frames of Marginal Effects from Regression Models. Journal of Open Source Software, 3(26), 772. 10.21105joss.00772

Lüdecke, D., Ben-Shachar, M., Patil, I., Waggoner, P., & Makowski, D. (2021). performance: An R Package for Assessment, Comparison and Testing of Statistical Models. Journal of Open Source Software, 6(60), 3139. 10.21105/joss.03139

Lundqvist, M., Miller, E. K., Nordmark, J., Liljefors, J., & Herman, P. (2024). Beta: Bursts of cognition. Trends in Cognitive Sciences, 28(7), 662–676. 10.1016/j.tics.2024.03.010

Maris, E., & Oostenveld, R. (2007). Nonparametric statistical testing of EEG- and MEG-data. Journal of Neuroscience Methods, 164(1), 177–190. 10.1016/j.jneumeth.2007.03.024

Mayer, E. A., Ryu, H. J., & Bhatt, R. R. (2023). The neurobiology of irritable bowel syndrome. Molecular Psychiatry, 28(4), 1451–1465. 10.1038/s41380-023-01972-w

Mehling, W. E., Acree, M., Stewart, A., Silas, J., & Jones, A. (2018). The Multidimensional Assessment of Interoceptive Awareness, Version 2 (MAIA-2). PloS One, 13(12), e0208034. 10.1371/journal.pone.0208034

Mihalik, A., Chapman, J., Adams, R. A., Winter, N. R., Ferreira, F. S., Shawe-Taylor, J., & Mourão-Miranda, J. (2022). Canonical Correlation Analysis and Partial Least Squares for Identifying Brain-Behavior Associations: A Tutorial and a Comparative Study. Biological Psychiatry: Cognitive Neuroscience and Neuroimaging, 7(11), 1055–1067. 10.1016/j.bpsc.2022.07.012

Minjoz, S., Jeanne, R., Pellissier, S., & Hot, P. (2026). Psychophysiological effects of heart rate variability biofeedback versus sham biofeedback: A randomized controlled trial. Biological Psychology, 206, 109254. 10.1016/j.biopsycho.2026.109254

Minjoz, S., Jeanne, R., Vercueil, L., Sabourdy, C., Sinniger, V., Bonaz, B., Hot, P., & Pellissier, S. (2025). Heart Rate Variability Biofeedback to Manage the Mental Health of Adults With Irritable Bowel Syndrome: A Pilot Study. Stress and Health, 41(1), e70015. 10.1002/smi.70015

Müller, S. J., Teckentrup, V., Rebollo, I., Hallschmid, M., & Kroemer, N. B. (2022). Vagus nerve stimulation increases gastric-brain coupling via a vagal afferent pathway. Brain Stimulation: Basic, Translational, and Clinical Research in Neuromodulation, 15(5), 1279–1289. 10.1016/j.brs.2022.08.019

Pascucci, D., Menétrey, M. Q., Passarotto, E., Luo, J., Paramento, M., & Rubega, M. (2025). EEG brain waves and alpha rhythms: Past, current and future direction. Neuroscience & Biobehavioral Reviews, 176, 106288. 10.1016/j.neubiorev.2025.106288

Pellissier, S., & Bonaz, B. (2017). The Place of Stress and Emotions in the Irritable Bowel Syndrome. In G. Litwack (Éd.), Vitamins and Hormones (Vol. 103, p. 327–354). Academic Press. 10.1016/bs.vh.2016.09.005

Plaisant, O., Courtois, R., Réveillère, C., Mendelsohn, G. A., & John, O. P. (2010). Validation par analyse factorielle du Big Five Inventory français (BFI-Fr). Analyse convergente avec le NEO-PI-R. Annales Médico-psychologiques, revue psychiatrique, 168(2), 97–106. 10.1016/j.amp.2009.09.003

Porciello, G., Monti, A., & Aglioti, S. M. (2018). How the stomach and the brain work together at rest. eLife, 7, e37009. 10.7554/eLife.37009

Porciello, G., Monti, A., Panasiti, M. S., & Aglioti, S. M. (2024). Ingestible pills reveal gastric correlates of emotions. eLife, 13, e85567. 10.7554/eLife.85567

Rebollo, I., Devauchelle, A.-D., Béranger, B., & Tallon-Baudry, C. (2018). Gastric-brain synchrony reveals a novel, delayed-connectivity resting-state network in humans. eLife, 7, e33321. 10.7554/eLife.33321

Rebollo, I., & Tallon-Baudry, C. (2022). The Sensory and Motor Components of the Cortical Hierarchy Are Coupled to the Rhythm of the Stomach during Rest. The Journal of Neuroscience, 42(11), 2205–2220. 10.1523/JNEUROSCI.1285-21.2021

Rebollo, I., Wolpert, N., & Tallon-Baudry, C. (2021). Brain-stomach coupling: Anatomy, functions, and future avenues of research. Current Opinion in Biomedical Engineering, 18, 100270. 10.1016/j.cobme.2021.100270

Richter, C. G., Babo-Rebelo, M., Schwartz, D., & Tallon-Baudry, C. (2017). Phase-amplitude coupling at the organism level: The amplitude of spontaneous alpha rhythm fluctuations varies with the phase of the infra-slow gastric basal rhythm. NeuroImage, 146, 951–958. 10.1016/j.neuroimage.2016.08.043

Sadowski, A., Dunlap, C., Lacombe, A., & Hanes, D. (2020). Alterations in Heart Rate Variability Associated With Irritable Bowel Syndrome or Inflammatory Bowel Disease: A Systematic Review and Meta-Analysis. Clinical and Translational Gastroenterology, 12(1), e00275. 10.14309/ctg.0000000000000275

Sequeira, H., Hot, P., Silvert, L., & Delplanque, S. (2009). Electrical autonomic correlates of emotion. International Journal of Psychophysiology: Official Journal of the International Organization of Psychophysiology, 71(1), 50–56. 10.1016/j.ijpsycho.2008.07.009

Seth, A. K. (2013). Interoceptive inference, emotion, and the embodied self. Trends in Cognitive Sciences, 17(11), 565–573. 10.1016/j.tics.2013.09.007

Spielberger, C. D., Bruchon-Schweitzer, M., & Paulhan, I. (1993). STAI-Y: Inventaire d’anxiété état-trait forme Y. Éditions du centre de psychologie appliquée.

Spielberger, C. D., Gorsuch, R., Lushene, R., Vagg, P., & Jacobs, G. (1983). Manual for the State-Trait Anxiety Inventory (Form Y1 - Y2). In Palo Alto, CA: Consulting Psychologists Press; IV.

Tort, A. B. L., Komorowski, R., Eichenbaum, H., & Kopell, N. (2010). Measuring Phase-Amplitude Coupling Between Neuronal Oscillations of Different Frequencies. Journal of Neurophysiology, 104(2), 1195–1210. 10.1152/jn.00106.2010

Vallat, R. (2018). Pingouin: Statistics in Python. Journal of Open Source Software, 3(31), 1026. 10.21105/joss.01026

Willem, C., Gandolphe, M.-C., Nandrino, J.-L., & Grynberg, D. (2022). French translation and validation of the Multidimensional Assessment of Interoceptive Awareness (MAIA-FR). Canadian Journal of Behavioural Science / Revue canadienne des sciences du comportement, 54(3), 234–240. 10.1037/cbs0000271

Wolpert, N., Rebollo, I., & Tallon-Baudry, C. (2020). Electrogastrography for psychophysiological research: Practical considerations, analysis pipeline, and normative data in a large sample. Psychophysiology, 57(9). 10.1111/psyp.13599

Yin, J., & Chen, J. D. Z. (2013). Electrogastrography: Methodology, Validation and Applications. Journal of Neurogastroenterology and Motility, 19(1), 5–17. 10.5056/jnm.2013.19.1.5

Ying-Chih, C., Yu-Chen, H., & Wei-Lieh, H. (2020). Heart rate variability in patients with somatic symptom disorders and functional somatic syndromes: A systematic review and meta-analysis. Neuroscience and Biobehavioral Reviews, 112, 336–344. 10.1016/j.neubiorev.2020.02.007

