## Supplementary Data for "Gastric-Brain Coupling Dynamics in Response to a Cognitive Challenge Reveal Distinct Autonomic and Psychological Correlates in Healthy Individuals and Patients with Irritable Bowel Syndrome Gastric-Brain Coupling Psychophysiological Correlates"

### Table 1. Cluster statistics for gastric-brain coupling in Healthy Adults and Adults With Irritable Bowel Syndrome.

| **Group** | **Frequency bands** | ***t_min_*** | ***t_mean_*** | ***t_max_*** | ***p_cluster_*** | ***N_electrodes_*** |
| --- | --- | --- | --- | --- | --- | --- |
| HA | delta | 2.06 | 2.61 | 3.93 | <.0001 | 47 |
|  | theta | 2.02 | 3.19 | 5.67 | <.0001 | 51 |
|  | alpha | 2.05 | 3.74 | 7.00 | <.0001 | 62 |
|  | beta | 2.04 | 3.48 | 7.83 | <.0001 | 57 |
|  | low gamma | 2.12 | 3.19 | 5.81 | <.0001 | 39 |
| IBS | delta | 2.97 | 3.35 | 3.61 | 0.0228 | 4 |
|  | delta | 2.14 | 2.75 | 4.48 | 0.001 | 14 |
|  | theta | 2.14 | 3.14 | 5.56 | 0.0001 | 44 |
|  | alpha | 2.09 | 2.68 | 3.84 | 0.0006 | 35 |
|  | beta | 2.17 | 3.03 | 5.57 | 0.0001 | 45 |
|  | low gamma | 2.08 | 2.65 | 4.36 | 0.0012 | 30 |
| IBS vs HA | delta | 2.15 | 2.33 | 2.64 | 0.6438 | 4 |
|  | theta | 6.68 | 7.81 | 8.37 | 0.2602 | 3 |
|  | alpha | 2.73 | 5.21 | 10.94 | 0.2077 | 6 |
|  | beta | 2.16 | 2.61 | 3.06 | 0.9213 | 2 |
|  | low gamma | 2.14 | 3.77 | 6.43 | 0.186 | 9 |

*Note. IBS, adults with Irritable Bowel Syndrome; HA, Healthy Adults. N_electrodes_ denotes the number of electrodes included in the cluster. Frequency bands were defined as follows: delta 1–4 Hz, theta 4–8 Hz, alpha 8–12 Hz, beta 12–30 Hz, and low gamma 30–45 Hz.*

### Table 2. Most Significant Cluster Characteristics for Post Hoc Contrasts of the Effect of Module on Gastric–Brain Coupling in Healthy Adults and Adults With Irritable Bowel Syndrome.

| **Group** | **Contrast** | ***t_mean_*** | ***t_min_*** | ***t_max_*** | ***p_cluster_*** | ***p_Holm_*** | ***N_electrodes_*** |
| --- | --- | --- | --- | --- | --- | --- | --- |
| HA | rest - task | 3.97 | 2.11 | 5.89 | 0.0001 | 0.0010 | 59 |
| HA | task - recovery010 | -2.98 | -5.19 | -2.17 | 0.0001 | 0.0010 | 41 |
| HA | task - recovery1020 | -3.43 | -4.96 | -2.05 | 0.0001 | 0.0010 | 57 |
| HA | task - recovery2030 | -3.88 | -5.21 | -2.03 | 0.0001 | 0.0010 | 58 |
| HA | recovery010 - recovery1020 | -2.21 | -2.45 | -2.06 | 0.0588 | 0.2352 | 7 |
| HA | recovery010 - recovery2030 | -2.64 | -3.63 | -2.03 | 0.0028 | 0.0168 | 38 |
| HA | recovery1020 - recovery2030 | -2.42 | -2.80 | -2.22 | 0.1625 | 0.3250 | 3 |
| HA | rest - recovery010 | 2.57 | 2.07 | 3.33 | 0.0093 | 0.0465 | 37 |
| HA | rest - recovery1020 | 2.42 | 2.08 | 2.88 | 0.0813 | 0.2439 | 6 |
| HA | rest - recovery2030 | 2.12 | 2.12 | 2.12 | 0.5578 | 0.5578 | 1 |
| IBS | rest - task | 4.32 | 2.58 | 6.37 | 0.0001 | 0.0010 | 63 |
| IBS | task - recovery010 | -3.24 | -5.38 | -2.10 | 0.0007 | 0.0042 | 47 |
| IBS | task - recovery1020 | -4.09 | -6.45 | -2.36 | 0.0001 | 0.0010 | 61 |
| IBS | task - recovery2030 | -4.72 | -6.75 | -2.23 | 0.0001 | 0.0010 | 63 |
| IBS | recovery010 - recovery1020 | -2.68 | -3.59 | -2.09 | 0.0040 | 0.0160 | 40 |
| IBS | recovery010 - recovery2030 | -3.22 | -4.48 | -2.24 | 0.0011 | 0.0055 | 40 |
| IBS | recovery1020 - recovery2030 |  |  |  |  |  |  |
| IBS | rest - recovery010 | 2.93 | 2.15 | 4.52 | 0.0004 | 0.0028 | 46 |
| IBS | rest - recovery1020 | 2.30 | 2.08 | 2.51 | 0.2844 | 0.8532 | 2 |
| IBS | rest - recovery2030 | 2.44 | 2.44 | 2.44 | 0.4315 | 0.8630 | 1 |
| IBS vs HA | rest |  |  |  |  |  |  |
| IBS vs HA | task | -0.27 | -0.93 | 0.46 | 0.1691 |  | 3 |
| IBS vs HA | recovery010 |  |  |  |  |  |  |
| IBS vs HA | recovery1020 | 1.11 | 0.58 | 1.78 | 0.1617 |  | 3 |
| IBS vs HA | recovery2030 | -0.26 | -0.26 | -0.26 | 0.4638 |  | 1 |

*Note. IBS, adults with Irritable Bowel Syndrome; HA, Healthy Adults. Recovery010 refers to the first 10 min of the recovery period, Recovery1020 to the second 10 min, and Recovery2030 to the third 10 min. Cells are shaded black when no significant cluster was identified. Cluster p values were corrected using the Holm–Bonferroni procedure across the family of pairwise module contrasts within each group. N_electrodes_ denotes the number of electrodes included in the cluster.*

### Mediations of gastric-brain coupling dynamics by EEG power

To determine whether the observed gastric–brain coupling effects were driven by changes in EEG power, within-subject mediation analyses were performed using the JSMediation package in R. Chance-corrected gastric–brain coupling values were log-transformed (real/chance ratio) to satisfy the normality and linearity assumptions of linear models. Mediation analyses were conducted separately for each electrode. Multiple-comparison correction was applied across the family of 64 electrodes using the Benjamini–Hochberg false discovery rate (FDR) procedure (*q* = .05). In these analyses, the total effect (path c) represents the effect of module on gastric–brain coupling. In these analyses, the total effect (path c) represents the effect of module on gastric–brain coupling. The mediation model tested whether this total effect could be decomposed into (i) an indirect effect of module on gastric–brain coupling through changes in EEG power and (ii) a remaining direct effect (path c′) of module on gastric–brain coupling after accounting for the indirect effect. The indirect effect is composed of path a (effect of module on EEG power) and path b (effect of EEG power on gastric–brain coupling while controlling for module) and is quantified as the product of these two paths (a × b). Following the joint-significance approach implemented in JSMediation, the indirect effect was considered significant only when both path a and path b were statistically significant. Significant mediation results summarized in this analysis showed that EEG power partially mediates gastric–brain coupling during recovery in frontal and parietal electrodes. In adults with IBS, EEG power partially mediates the gastric–brain coupling response to the task in frontal and parietal electrodes, but no significant mediation was observed during recovery.

Because mediation analyses were performed separately for each electrode, Table 5 provides a summary of the significant findings by reporting the median estimate, standard error, t value, and p value across electrodes surviving FDR correction (*p_corrected_* < .05) for a given effect. Figure 1 displays the topographical distributions of the indirect, direct, and total effect estimates in HA and adults with IBS for contrasts in which a significant indirect effect was observed in at least one group.

This analysis showed that EEG power partially mediates gastric–brain coupling during recovery in frontal and parietal electrodes. In adults with IBS, EEG power partially mediates the gastric–brain coupling response to the task in frontal and parietal electrodes, but no significant mediation was observed during recovery.

### Table 3. Statistical Results of Within-Subject Mediation Analyses of Gastric–Brain Coupling by EEG Power in Healthy Adults and Adults With Irritable Bowel Syndrome

| **Group** | **Contrast** | **Effect** | **Path** | | ***Estimate*** | ***SE*** | ***t*** | ***p_FDR_*** | ***N_electrodes_*** |
| --- | --- | --- | --- | --- | --- | --- | --- | --- | --- |
| HA | task - recovery010 | Indirect | a | -0.28 | | 0.07 | -4.15 | 0.0001 | 1 |
| HA |  |  | b | 0.63 | | 0.15 | 4.09 | 0.0002 | 1 |
| HA | task - recovery1020 | Indirect | a | -0.29 | | 0.06 | -4.59 | <0.0001 | 20 |
| HA |  |  | b | 0.67 | | 0.20 | 3.19 | 0.0027 | 20 |
| HA | task - recovery2030 | Indirect | a | -0.28 | | 0.06 | -5.33 | 0.0001 | 2 |
| HA |  |  | b | 0.74 | | 0.19 | 3.99 | 0.0003 | 2 |
| HA | task - recovery010 | Total | c | -0.22 | | 0.07 | -3.35 | 0.0017 | 47 |
| HA | task - recovery1020 | Total | c | -0.29 | | 0.07 | -4.21 | 0.0001 | 60 |
| HA | task - recovery2030 | Total | c | -0.35 | | 0.07 | -4.69 | 0.0000 | 63 |
| HA | recovery 010 - recovery2030 | Total | c | -0.16 | | 0.05 | -3.34 | 0.0017 | 52 |
| HA | recovery 1020 - recovery2030 | Total | c | -0.13 | | 0.04 | -3.69 | 0.0008 | 4 |
| HA | rest - task | Total | c | -0.35 | | 0.07 | -5.02 | <0.0001 | 63 |
| HA | rest - recovery010 | Total | c | -0.17 | | 0.06 | -3.07 | 0.0037 | 49 |
| HA | task - recovery010 | Direct | c' | -0.27 | | 0.07 | -3.37 | 0.0016 | 19 |
| HA | task - recovery1020 | Direct | c' | -0.26 | | 0.08 | -3.34 | 0.0018 | 16 |
| HA | task - recovery2030 | Direct | c' | -0.30 | | 0.09 | -3.15 | 0.0030 | 42 |
| HA | recovery 010 - recovery2030 | Direct | c' | -0.15 | | 0.05 | -3.17 | 0.0029 | 47 |
| HA | recovery 1020 - recovery2030 | Direct | c' | -0.13 | | 0.04 | -3.72 | 0.0007 | 4 |
| HA | rest - task | Direct | c' | -0.31 | | 0.10 | -3.35 | 0.0017 | 44 |
| HA | rest - recovery010 | Direct | c' | -0.16 | | 0.06 | -2.86 | 0.0065 | 38 |
| IBS | rest - task | Indirect | a | -0.53 | | 0.12 | -4.70 | 0.0001 | 31 |
| IBS | rest - task |  | b | 0.44 | | 0.15 | 2.93 | 0.0079 | 31 |
| IBS | task - recovery010 | Total | c | -0.27 | | 0.08 | -3.34 | 0.0029 | 56 |
| IBS | task - recovery1020 | Total | c | -0.40 | | 0.09 | -4.67 | 0.0001 | 63 |
| IBS | task - recovery2030 | Total | c | -0.45 | | 0.08 | -5.58 | <0.0001 | 63 |
| IBS | recovery 010 - recovery1020 | Total | c | -0.20 | | 0.07 | -2.90 | 0.0081 | 26 |
| IBS | recovery 010 - recovery2030 | Total | c | -0.23 | | 0.07 | -3.62 | 0.0014 | 40 |
| IBS | rest - task | Total | c | -0.44 | | 0.08 | -4.97 | 0.0001 | 64 |
| IBS | rest - recovery010 | Total | c | -0.20 | | 0.07 | -2.90 | 0.0081 | 50 |
| IBS | task - recovery010 | Direct | c' | -0.35 | | 0.11 | -3.39 | 0.0028 | 9 |
| IBS | task - recovery1020 | Direct | c' | -0.39 | | 0.12 | -3.31 | 0.0034 | 50 |
| IBS | task - recovery2030 | Direct | c' | -0.44 | | 0.11 | -4.05 | 0.0006 | 50 |
| IBS | recovery 010 - recovery1020 | Direct | c' | -0.26 | | 0.07 | -3.87 | 0.0009 | 2 |
| IBS | recovery 010 - recovery2030 | Direct | c' | -0.27 | | 0.07 | -3.99 | 0.0007 | 42 |
| IBS | rest - task | Direct | c' | -0.37 | | 0.10 | -3.43 | 0.0025 | 23 |
| IBS | rest - recovery010 | Direct | c' | -0.22 | | 0.07 | -2.73 | 0.0126 | 32 |

*Note. SE, standard error; IBS, adults with Irritable Bowel Syndrome; HA, Healthy Adults. recovery010 refers to the first 10 min of the recovery period, recovery1020 to the second 10 min, and recovery2030 to the third 10 min. Shaded cells highlight the significant indirect effects.*

### Figure 1. Topographical representation of within-subject mediation estimates for gastric–brain coupling contrasts in Healthy Adults (HA) and adults with Irritable Bowel Syndrome (IBS).

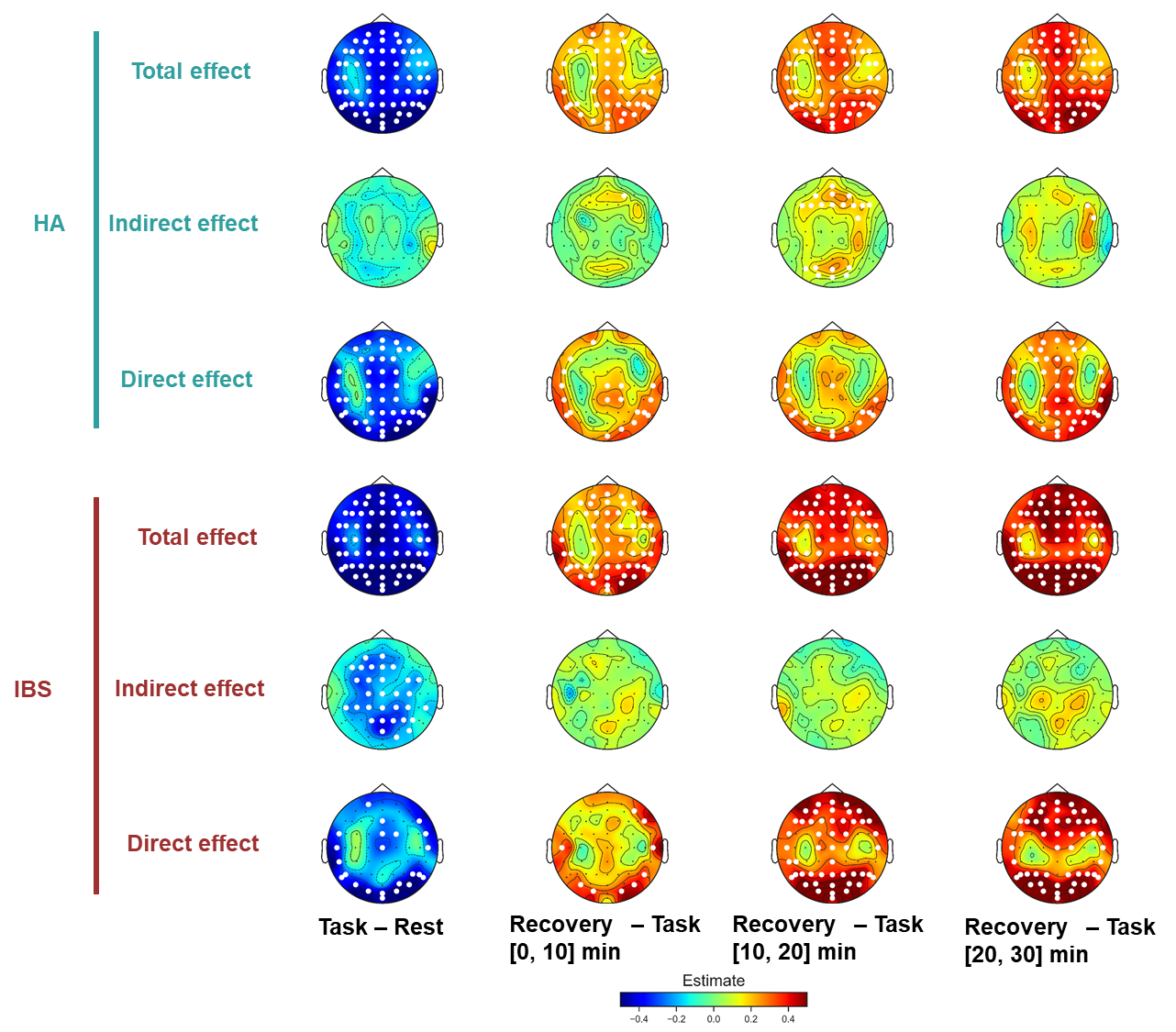

*Note. IBS, adults with Irritable Bowel Syndrome; HA, Healthy Adults. Electrodes showing significant effects (FDR-corrected) are highlighted in white.*

### Table 4. Statistical Results of Linear Mixed-Effects Models by Outcome Variable.

| **Variable** | **Effect** | $\chi^{2}$ | ***df*** | ***p*** | $\eta_{p}^{2}$ |
| --- | --- | --- | --- | --- | --- |
| EDA-SCR | (Intercept) | 299.81 | 1 | <.001 |  |
|  | module | 206.76 | 4 | <.001 | 0.250 |
|  | session | 2.46 | 1 | 0.117 | 0.004 |
|  | group | 0.32 | 1 | 0.573 | 0.005 |
|  | module:session | 0.16 | 4 | 0.997 | <.001 |
|  | module:group | 3.93 | 4 | 0.416 | 0.006 |
|  | session:group | 0.00 | 1 | 0.952 | <.001 |
|  | module:session:group | 0.88 | 4 | 0.928 | 0.001 |
| HRV-RMSSD | (Intercept) | 3517.44 | 1 | <.001 |  |
|  | module | 17.29 | 4 | 0.002 | 0.027 |
|  | session | 0.00 | 1 | 0.964 | <.001 |
|  | group | 5.98 | 1 | 0.014 | 0.080 |
|  | module:session | 1.71 | 4 | 0.789 | 0.003 |
|  | module:group | 4.11 | 4 | 0.392 | 0.007 |
|  | session:group | 7.54 | 1 | 0.006 | 0.012 |
|  | module:session:group | 1.81 | 4 | 0.770 | 0.003 |
| Gastric slow wave frequency | (Intercept) | 10796.61 | 1 | <.001 |  |
|  | module | 1.37 | 4 | 0.849 | 0.002 |
|  | session | 1.82 | 1 | 0.177 | 0.003 |
|  | group | 0.86 | 1 | 0.355 | 0.012 |
|  | module:session | 1.60 | 4 | 0.809 | 0.003 |
|  | module:group | 3.20 | 4 | 0.525 | 0.005 |
|  | session:group | 0.08 | 1 | 0.774 | <.001 |
|  | module:session:group | 1.39 | 4 | 0.845 | 0.002 |
| Gastric slow wave standard deviation of cycle duration | (Intercept) | 861.31 | 1 | <.001 |  |
|  | module | 7.37 | 4 | 0.118 | 0.012 |
|  | session | 3.82 | 1 | 0.051 | 0.006 |
|  | group | 26.13 | 1 | <.001 | 0.275 |
|  | module:session | 1.90 | 4 | 0.755 | 0.003 |
|  | module:group | 8.93 | 4 | 0.063 | 0.014 |
|  | session:group | 0.54 | 1 | 0.464 | 0.001 |
|  | module:session:group | 0.87 | 4 | 0.928 | 0.001 |
| Gastric slow wave power | (Intercept) | 2857.52 | 1 | <.001 |  |
|  | module | 4.76 | 4 | 0.312 | 0.008 |
|  | session | 0.03 | 1 | 0.855 | <.001 |
|  | group | 2.39 | 1 | 0.122 | 0.034 |
|  | module:session | 0.54 | 4 | 0.970 | 0.001 |
|  | module:group | 3.17 | 4 | 0.529 | 0.005 |
|  | session:group | 0.34 | 1 | 0.559 | 0.001 |
|  | module:session:group | 2.70 | 4 | 0.609 | 0.004 |
| Perceived stress | (Intercept) | 359.30 | 1 | <.001 |  |
|  | module | 56.77 | 2 | <.001 | 0.175 |
|  | session | 2.67 | 1 | 0.102 | 0.002 |
|  | group | 1.05 | 1 | 0.304 | <.001 |
|  | module:session | 6.10 | 2 | 0.047 | 0.009 |
|  | module:group | 6.97 | 2 | 0.031 | 0.022 |
|  | session:group | 3.06 | 1 | 0.080 | 0.005 |
|  | module:session:group | 1.96 | 2 | 0.376 | 0.006 |

*Note. Linear mixed-effects models (LMMs) were fitted to square-root-transformed EDA-SCR, log-transformed HRV-RMSSD, log-transformed gastric slow-wave power, standard deviation of slow-wave cycle duration, and square-root-transformed perceived stress scores to satisfy model assumptions. df, degrees of freedom.*

### Table 5. Post Hoc Contrasts for Significant Module Effects and Module × Group Interactions.

| **Post-hoc tables EDA-SCR** | | | | | |
| --- | --- | --- | --- | --- | --- |
| **Contrast** | **Estimate** | ***SE*** | ***t*** | ***df*** | ***p*** |
| rest - task | -0.21 | 0.02 | -9.89 | 638.54 | <.001 |
| rest - recovery010 | -0.25 | 0.02 | -11.29 | 638.54 | <.001 |
| rest - recovery1020 | -0.04 | 0.02 | -2.05 | 638.57 | 0.243 |
| rest - recovery2030 | -0.07 | 0.02 | -3.11 | 638.54 | 0.017 |
| task - recovery010 | -0.03 | 0.02 | -1.40 | 638.54 | 0.628 |
| task - recovery1020 | 0.17 | 0.02 | 7.83 | 638.57 | <.001 |
| task - recovery2030 | 0.15 | 0.02 | 6.78 | 638.54 | <.001 |
| recovery010 - recovery1020 | 0.20 | 0.02 | 9.23 | 638.57 | <.001 |
| recovery010 - recovery2030 | 0.18 | 0.02 | 8.18 | 638.54 | <.001 |
| recovery1020 - recovery2030 | -0.02 | 0.02 | -1.06 | 638.57 | 0.829 |
| **Post-hoc tables HRV-RMSSD** | | | | | |
| **Contrast** | **Estimate** | ***SE*** | ***t*** | ***df*** | ***p*** |
| rest - task | 0.05 | 0.02 | 2.16 | 639.54 | 0.196 |
| rest - recovery010 | -0.04 | 0.02 | -1.47 | 639.54 | 0.581 |
| rest - recovery1020 | -0.02 | 0.02 | -0.96 | 639.54 | 0.873 |
| rest - recovery2030 | -0.02 | 0.02 | -0.97 | 639.54 | 0.868 |
| task - recovery010 | -0.09 | 0.02 | -3.63 | 639.54 | 0.003 |
| task - recovery1020 | -0.08 | 0.02 | -3.12 | 639.54 | 0.016 |
| task - recovery2030 | -0.08 | 0.02 | -3.13 | 639.54 | 0.016 |
| recovery010 - recovery1020 | 0.01 | 0.02 | 0.51 | 639.54 | 0.986 |
| recovery010 - recovery2030 | 0.01 | 0.02 | 0.50 | 639.54 | 0.987 |
| recovery1020 - recovery2030 | 0.00 | 0.02 | -0.01 | 639.54 | 1.000 |
| **Post-hoc tables perceived stress** | | | | | |
| **Contrast** | **Estimate** | ***SE*** | ***t*** | ***df*** | ***p*** |
| rest - task | -0.21 | 0.05 | -4.10 | 325.38 | <.001 |
| rest - recovery010 | 0.20 | 0.05 | 3.96 | 325.38 | <.001 |
| task - recovery010 | 0.41 | 0.05 | 8.06 | 325.38 | <.001 |
| linear: [IBS-HA] | 0.08 | 0.10 | 0.78 | 325.38 | 0.438 |
| quadratic: [IBS-HA] | 0.45 | 0.18 | 2.52 | 325.38 | 0.012 |

*Note. SE, standard error; df, degrees of freedom; IBS, adults with irritable bowel syndrome; HA, Healthy Adults. Pairwise contrast p values were corrected using the Tukey method. Recovery010 refers to the first 10 min of the recovery period, Recovery1020 to the second 10 min, and Recovery2030 to the third 10 min. Linear: [IBS–HA] and quadratic: [IBS–HA] refer to the interaction contrasts testing the linear and quadratic effects of module between the IBS and HA groups, respectively.*

### Table 6. Group Differences in Psychological Traits Between Adults With Irritable Bowel Syndrome and Healthy Adults*.*

| **Variable** | **Descriptive IBS** | **Descriptive HA** | **test** | **statistic** | ***p_uncorrected_*** | ***p_FDR_*** | ***Effect size*** |
| --- | --- | --- | --- | --- | --- | --- | --- |
| STAIYB | 51.3 ± 11.9 | 46.1 ± 11.3 | t-test | 1.73 | 0.088 | 0.280 | 0.45 |
| BFI_N | 28.0 ± 7.9 | 23.5 ± 7.8 | t-test | 2.23 | 0.029 | 0.147 | 0.57 |
| MAIA-2 Noticing | 3.3 ± 0.8 | 3.3 ± 0.8 | t-test | -0.40 | 0.691 | 0.817 | 0.10 |
| MAIA-2 Not-distracting | 2.2 (1.0) | 2.2 (1.5) | Mann-Whitney | 385.50 | 0.112 | 0.280 | 0.24 |
| MAIA-2 Not-worrying | 2.8 (1.0) | 2.6 (1.2) | Mann-Whitney | 488.00 | 0.817 | 0.817 | 0.04 |
| MAIA-2 Attention regulation | 2.4 ± 0.9 | 2.5 ± 0.9 | t-test | -0.37 | 0.710 | 0.817 | 0.10 |
| MAIA-2 Emotional awareness | 3.4 (1.2) | 3.7 (1.4) | Mann-Whitney | 482.00 | 0.756 | 0.817 | 0.05 |
| MAIA-2 Self-regulation | 2.2 ± 1.0 | 2.4 ± 1.1 | t-test | -0.49 | 0.627 | 0.817 | 0.13 |
| MAIA-2 Body listening | 2.1 ± 1.3 | 2.2 ± 1.3 | t-test | -0.41 | 0.681 | 0.817 | 0.11 |
| MAIA-2 Trusting | 2.0 (1.8) | 3.3 (1.4) | Mann-Whitney | 326.50 | 0.018 | 0.147 | 0.35 |

*Note. Descriptive statistics are reported as mean ± standard deviation for normally distributed variables and median (interquartile range) for non-normally distributed variables. Accordingly, parametric between-group comparisons were performed using independent-samples t-tests, whereas non-parametric comparisons were performed using Mann–Whitney U tests. Reported effect sizes include Cohen’s d for parametric comparisons and rank-biserial correlation for non-parametric comparisons.*
